# Anatomical organization and origins of VGLUT3-positive axon terminals in the lateral septum

**DOI:** 10.64898/2026.06.10.729142

**Authors:** Lina I. Elvers, Suzanne van der Veldt, Justine Fortin-Houde, Guillaume Ducharme, Bénédicte Amilhon

**Author notes:** Corresponding author: Bénédicte Amilhon, Université de Montréal, Département de neurosciences, CHU Sainte-Justine Azrieli Research Center, Montréal (QC) Canada.

## Abstract

The lateral septum (LS) integrates afferents from multiple brain regions, including the raphe nuclei. The organization of these inputs contributes to the regionalization of LS functions, for example spatial coding in dorsal LS and emotional regulation in ventral LS. Raphe-LS projections include glutamatergic axons expressing the vesicular glutamate transporter type 3 (VGLUT3), which often form pericellular baskets around LS neurons. This study provides an anatomical characterization of the organization and origins of VGLUT3-positive (VGLUT3^+^) raphe inputs to the LS. We mapped VGLUT3^+^ axon terminal density across the rostro-caudal extent of the LS and quantified colocalization with serotonin (5-HT) using immunohistochemistry. Our results showed that VGLUT3 density was highest in the ventral LS, whereas VGLUT3/5-HT colocalization was strongest in the dorsal LS. Retrograde viral vector-mediated tracing identified predominant inputs from the median raphe and B9 neuron group. Interestingly, the ventral hippocampus, a functionally related region which is known to also receive raphe VGLUT3 inputs, showed collaterals with the LS. Additional VGLUT3^+^ inputs to the LS arose from the interpeduncular nucleus, bed nucleus of the stria terminalis, nucleus incertus and pontine central gray. Anterograde tracing revealed that inputs from these brain regions target distinct and largely non-overlapping domains in the LS. Our findings highlight multiple sources of VGLUT3^+^ inputs to the LS, beyond the raphe nuclei, and suggest that distinct VGLUT3 circuits could contribute to LS functional specialization.

## Introduction

The lateral septum (LS) is a central limbic structure that integrates hippocampal, hypothalamic, amygdala, and brainstem inputs to regulate emotional, motivational, and social behaviors (Besnard & Leroy, 2022; Menon et al., 2022; Patel, 2022; Risold & Swanson, 1997; Rizzi-Wise & Wang, 2021; Sheehan et al., 2004; Swanson & Cowan, 1979; Wirtshafter & Wilson, 2021; Yeates et al., 2022). Early anatomical studies revealed that the LS is not a homogeneous structure but rather consists of dorsal (dLS), intermediate (iLS), and ventral (vLS) subunits, each embedded within distinct functional circuits (Risold & Swanson, 1997a; Risold & Swanson, 1997b; Swanson & Cowan, 1979). Recent work has emphasized that these subdivisions perform highly specific roles in shaping adaptive and maladaptive behaviors: the dLS has been implicated in spatial representations, defensive response, threat detection, and social behavior, whereas the vLS participates in anxiety regulation, motivational processes, and appetitive behaviors (Besnard et al., 2019; Besnard & Leroy, 2022; Leutgeb & Mizumori, 2002; Menon et al., 2022; Tingley & Buzsáki, 2018; van der Veldt et al., 2021; Wirtshafter & Wilson, 2019; Yeates et al., 2022). LS circuits mediate interactions between hippocampal contextual information and hypothalamic or midbrain action-selection pathways, acting as a critical hub for various functions such as locomotion, motivation and feeding (Patel, 2022; van der Veldt et al., 2021; Wirtshafter & Wilson, 2019; Wirtshafter & Wilson, 2020; Wirtshafter & Wilson, 2021).

A defining feature of the LS is its rich neuromodulatory innervation. Serotonergic (5-HT) afferents from the dorsal and median raphe nuclei densely innervate the entire LS (Hale & Lowry, 2011; Lowry et al., 2005; Muzerelle et al., 2016). Serotonergic raphe neurons themselves are highly heterogeneous, comprising distinct developmental lineages and functionally specialized subgroups organized along the rostro-caudal and dorso-ventral axes of the raphe nuclei (Alonso et al., 2013; Commons, 2016; Muzerelle et al., 2016; Senft et al., 2021). Within serotonergic systems, the vesicular glutamate transporter type 3 (VGLUT3 or Slc17a8) has emerged as a key factor in 5-HT neuron diversity. VGLUT3 is atypical in that it can be expressed in neurons that typically use a primary transmitter other than glutamate, including serotonergic, cholinergic, and GABAergic neurons (Gras et al., 2002; Herzog et al., 2004; Schafer et al., 2002). Through a mechanism known as vesicular-filling synergy, VGLUT3 enhances vesicular content of acetylcholine and monoamines, thereby increasing the release of neurotransmitters (Amilhon et al., 2010; Cristofari et al., 2022; Favier et al., 2021; Gras et al., 2008).

VGLUT3^+^ afferents form a prominent component of LS innervation. In rats and mice, VGLUT3 terminals form characteristic pericellular baskets (PCBs) around the somata and proximal dendrites of LS neurons, suggesting a potent modulatory influence on local microcircuits (Riedel et al., 2008). Although a subset of VGLUT3^+^ inputs to the LS have been shown to arise from the raphe nuclei (Senft & Dymecki, 2021; Senft et al., 2021), a comprehensive overview of the sources of VGLUT3 terminals, their topographical organization and colocalization with 5-HT across the entire LS is lacking. Functionally, the VGLUT3 system has been implicated in anxiety, stress reactivity, fear generalization, and emotional memory, as deficiency or dysfunction of VGLUT3 leads to impairments in these behaviors (Amilhon et al., 2010; de Almeida et al., 2023; Henderson et al., 2024; Pujol et al., 2023). Given the established roles of both the LS and the raphe nuclei in affective regulation, VGLUT3-dependent glutamatergic signaling may represent an important mechanism through which these circuits modulate affective states.

A subset of VGLUT3^+^ axon terminals in the LS may also arise from other sources than the raphe nuclei. The LS receives dense inputs from the bed nucleus of stria terminalis (BNST), a region that contains VGLUT3-expressing neurons (Herzog et al., 2004; Schafer et al., 2002). In addition, the LS also receives projections from multiple brainstem regions, including the nucleus incertus (NI) and pontine central gray (PCG), which contain sparse and poorly characterized populations of VGLUT3^+^ neurons (see *Slc17a8* expression in the Allen Institute in situ hybridization atlas) (Dong & Swanson, 2004; Goto et al., 2001; Rizzi-Wise & Wang, 2021). Taken together, the precise origins and topographical organization of VGLUT3^+^ axon terminals in the LS, their extent of colocalization with 5-HT, and their potential collateralization to functionally related regions such as the ventral hippocampus await clarification.

The present study addresses these gaps by providing an anatomical characterization of VGLUT3^+^ inputs to the LS in mice, using immunohistochemistry and viral circuit-tracing approaches. Our results showed that VGLUT3+ axon terminals are topographically organized along the rostro-caudal and dorso-ventral axes of the LS. In particular, the vLS displays the highest density of VGLUT3+ terminals but low levels of colocalization with 5-HT; whereas the dLS shows the opposite pattern. Retrograde tracing identified LS-projecting VGLUT3 neurons (VGLUT3^LS^) in multiple regions: the ventral sub-divisions of the raphe nuclei (MRR and B9), the interpeduncular nucleus (IP), the BNST, the NI and the PCG. Axon terminals arising from these regions had varied morphology, with NI and PCG sending thin varicosed axons while IP, raphe and BNST VGLUT3 terminals formed PCBs, and innervated distinct LS territories. Together, these results provide a comprehensive anatomical framework for understanding how VGLUT3-dependent glutamatergic and glutamate/5-HT co-transmission are organized within LS circuits. By defining the topography, molecular phenotype, and origins of VGLUT3 inputs to distinct LS subdivisions, this study reveals previously uncharacterized features of septal organization and identifies candidate pathways through which VGLUT3 signaling may influence affective, motivational, and stress-related behaviors.

## Materials and methods

All procedures were approved by the Comité Institutionnel de Bonne Pratique Animale en Recherche (CIBPAR) at the CHU Sainte-Justine Research Center, in accordance with the standards of the Conseil Canadien de Protection des animaux (CCPA). Mice were housed under a 12-hour light/dark cycle with *ad libitum* access to food and water. For this study, we used adult male and female heterozygotes VGLUT3-Cre mice maintained on a C57BL/6J (Jackson Laboratory, #000664) genetic background (ages between 2.5 and 4 months) and wild-type (WT) littermates. VGLUT3-Cre strain is a gift from Dr Salah El Mestikawy (The Douglas Research Center, McGill University, Montréal). Specificity of Cre expression in the raphe was established in a previous study (Fortin-Houde et al., 2023).

### Viral injections

For all surgeries, mice were anesthetized with 5% isoflurane prior to virus injections and maintained under anesthesia with 1.25% isoflurane. Body temperature was maintained at 37 °C with a heating pad; mice received 0.3 mL of carprofen intraperitoneally before the start of the surgery.

### Retrograde tracing experiment

To label neurons in the raphe nuclei that project either to the LS or the vHP, we injected n = 12 VGLUT3-Cre mice (7 males, 4 females and 1 WT control for viral specificity) with the viral vectors AAV2retro-Ef1α-DIO-eYFP (5.5×10^12^ GC/ml, Canadian Neurophotonics Vectorology Core, RRID:SCR_016477, ULaval) and AAV2retro-CAG-FLEX-tdTomato (2.1×10^13^ GC/ml provided by Edward Boyden, catalog #28306-AAVrg, Addgene) in the LS and vHP, respectively. n = 2 additional mice were injected in the LS only, with the viral vector AAV2retro-Ef1α-DIO-eYFP. Stereotaxic injections (David Kopf Instruments) were done bilaterally with eYFP-expressing virus injected in the LS (50 nL per hemisphere) and tdTomato-expresing virus in the vHP (200 nL per hemisphere). The following coordinates were used: LS: anteroposterior (AP): +0.50 mm; mediolateral (ML): ±0.60 mm; dorsoventral (DV): -3.70 mm; vHP: AP: -3.64 mm; ML: ±3.52 mm; DV: -4.22 mm, with a 7° lateral angle.

### Anterograde tracing experiment

The origins of VGLUT3^+^ inputs to the lateral septum (LS) were mapped using anterograde tracing with AAVdj-EF1a-DIO-eYFP (9.0×10^12^ GC/ml, Canadian Neurophotonics Vectorology Core) and AAV2/9-hSyn-DIO-mCherry (7.0×10^12^ GC/ml, Canadian Neurophotonics Vectorology Core). n = 11 mice (5 males, 5 females and 1 male WT for viral specificity control) were injected in the BNST (coordinates: AP: 0.62 mm, ML: ±0.56 mm, DV: -4.40 mm, unilateral injection of 10-25 nL of AAVdj-EF1a-DIO-eYFP viral vector), the median raphe nucleus and interpeduncular nucleus (MnR/IP, AP: -4.03 mm, ML: ±0.83 mm, DV: -4.73 mm, 10° lateral angle, injection of 10-150 nL of either viral vector) and/or the nucleus incertus and pontine central gray (NI/ PCG, AP: -5.41 mm, ML: 0 mm, DV: -4.20 mm, injection of 10-50 nL of AAVdj-EF1a-DIO-eYFP viral vector).

### Tissue preparation

Five weeks post-injection, mice were perfused transcardially with 4% paraformaldehyde (PFA) in Phosphate Buffer Saline (PBS), following anesthesia using Ketamine (80 mg/kg), Xylazine (12.5 mg/kg) and Acepromazine (2.5 mg/kg), injected intraperitoneally. The brains were kept in 4% PFA for 48 h at 4°C. Brains were cut into 40 µm (VGLUT3^+^ and 5-HT^+^ boutons quantification) or 50 µm coronal sections (retrograde and anterograde experiments), using a vibratome (Leica BioSystems, VT1200).

### Immunofluorescence

#### VGLUT3^+^ and 5-HT^+^ boutons in the lateral septum

To identify VGLUT3^+^ and 5-HT^+^ terminals in the LS, a double immunostaining was performed on n = 12 WT mice (6 males, 6 females). Brain sections were incubated for 48 h at 4°C with the following primary antibodies: rabbit anti-VGLUT3 (1:1000, catalog #135203, Synaptic Systems) and goat anti-5-HT (1:2000, catalog #ab66047, Abcam), following 3 x 45 min of blocking using PBS-gelatin 0.45%-triton 0.25% solution (PGT). The sections were then washed 3 x 15 min in PGT followed by 1.5 h incubation in the following secondary antibodies: donkey anti-rabbit-A488 (1:2000, catalog # A21206, Life Technologies) and donkey anti-goat-A555 (1:2000, catalog #A21432, Life Technologies). Sections were washed 3 x 15 min in PBS and mounted on glass slides using Fluoromount-G™ Mounting Medium, with DAPI (catalog #00-4959-52, Invitrogen).

#### Tracing experiments

Using the same protocol, sections were incubated in the following primary antibodies: chicken anti-Green Fluorescent Protein (GFP, 1:2000, catalog #A10262, Invitrogen), goat anti-Red Fluorescent Protein (RFP, 1:10 000, catalog #200-101-379, VWR (Rockland)) and either rabbit anti-Tryptophan Hydroxylase 2 (Tph2, 1:2000, catalog #NB100-74555, Novus Biological) or rabbit anti-VGLUT3 (as described previously). Secondary antibodies used were the following: donkey anti-goat-A555 (as described previously), donkey anti-chicken-A488 (1:2000, catalog #703-545-155, Jackson Immunoresearch) and donkey anti-rabbit-A647 (1:2000, catalog #A31573, Life technologies). Additional ChAT (choline acetyl transferase) immunostaining was performed on some of the sections, using a goat anti-ChAT antibody (1:1000, catalog #AB144P, Millipore) and donkey anti-goat-A555. Relaxin-3 (Rln3) immunostaining was used to delineate the nucleus incertus, using a rabbit anti-Rln3 antibody (1:100, catalog #26075-1-AP, Thermofisher) and donkey anti-rabbit-A555.

### Image acquisition and quantification

#### VGLUT3^+^ and 5-HT^+^ boutons in the lateral septum

Images in Figure 1 (VGLUT3 expression pattern in the LS), were acquired using an inverted wide-field fluorescence microscope (Leica DMi8) with 10x magnification. For the quantification of VGLUT3^+^ puncta density in the LS and colocalization with 5-HT, images were acquired using a laser scanning confocal microscope on an inverted stand (Leica TCS SP8-DLS). Acquisition settings included a resolution of 2048 × 2048 pixels, a zoom factor of 2.0, and a scanning speed of 600 Hz. Imaging was performed at 20× magnification. For each LS level (120 µm spacing between levels), a single focal plane was acquired. The density of VGLUT3^+^ punctas and colocalization with 5-HT were determined using the FIJI plugin ComDet (Katrukha, 2020; Schindelin et al., 2012). Rectangular ROIs were drawn to encompass the lateral septum (LS), and areas outside the LS were manually excluded. Within the ROIs, puncta were detected independently in each channel (VGLUT3 and 5-HT), based on user-defined thresholds. VGLUT3^+^ and 5-HT^+^ punctas were considered colocalized if the distance between center of detected area was inferior to 2 pixels. The coordinates of detected puncta were used to quantify the density of VGLUT3^+^ puncta and proportion of these puncta colocalizing with 5-HT using custom MATLAB scripts. VGLUT3^+^ puncta density in dLS and vLS sub-regions was calculated in hand-delimitated ROIs to quantify total VGLUT3^+^ surface occupancy for each subregion (dLS or vLS) and proportion of colocalization with 5-HT. LS borders and subregions were delimitated based on VGLUT3 staining (delineating the nucleus accumbens and BNST) and *The mouse brain atlas* by K. B.J. Paxinos and G. Franklin (Paxinos & Franklin, 2019). The reliability of the FIJI plugin ComDet was evaluated by comparing automated puncta detection with manual quantification within a defined ROI from one section per animal. This comparison indicated that ComDet produced a slight (5.67 ± 7,06%) overestimation of puncta counts (analysis not shown).

**Figure 1.**
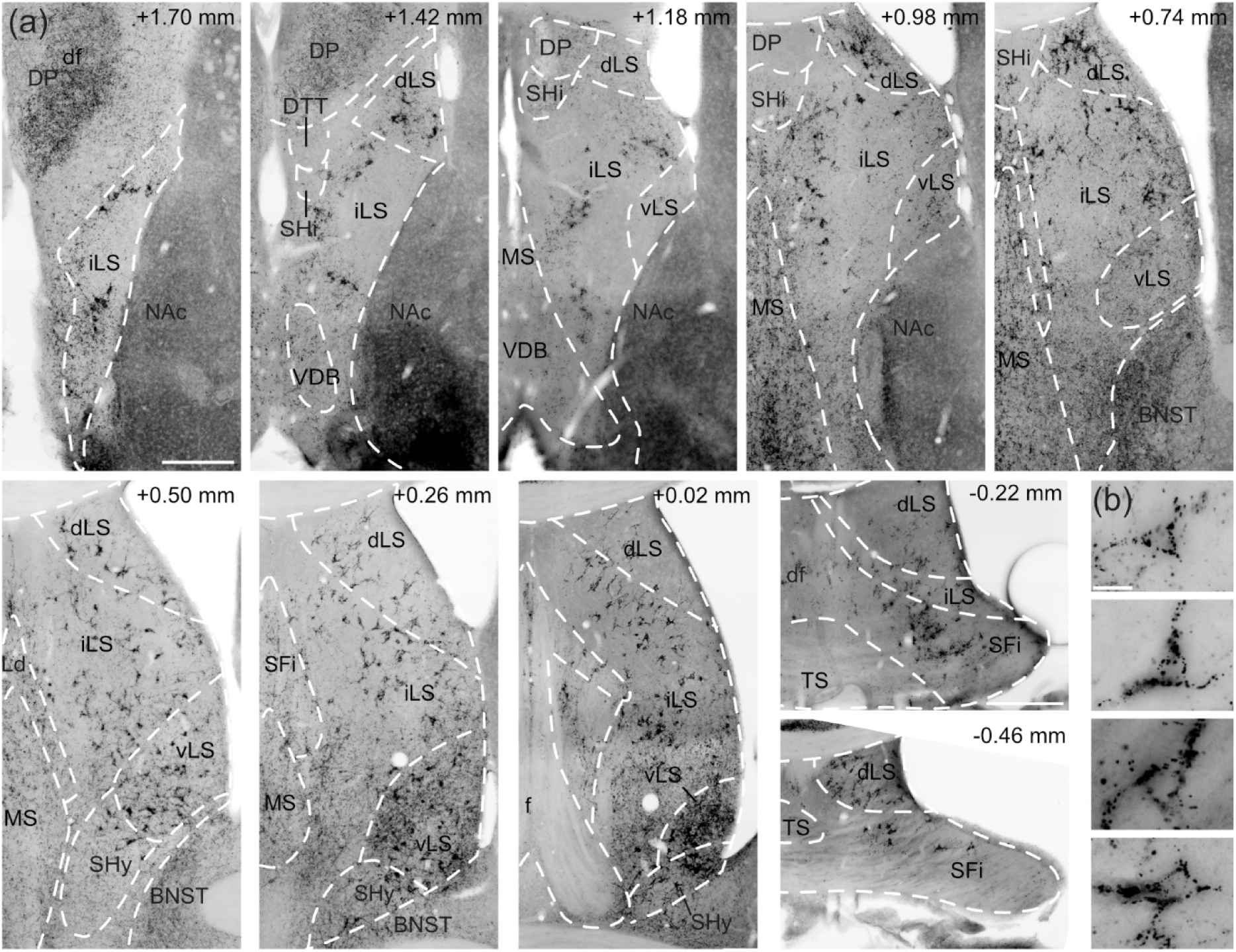
Distribution of VGLUT3^+^ axon terminals in the LS. (a) Immunohistochemistry showing VGLUT3^+^ axon terminal distribution at various rostro-caudal LS levels. Distance from bregma is indicated in the top right corner for each level. (b) Examples of VGLUT3^+^ axon terminals forming pericellular baskets (PCBs) in the LS. All subdivisions of the LS were determined according to *The mouse brain, K. B.J. Paxinos and G. Franklin (Paxinos & Franklin, 2019)*. Scale bar 300 µm (+1.70 to +0.02 mm), 400 µm (-0.22, -0.46 mm), 20 µm (b). The images were taken from n = 1 WT female mouse. All LS related regions are highlighted in white initials. BNST: bed nucleus of stria terminalis; df: dorsal fornix; dLS: lateral septal nucleus, dorsal part; DP: dorsal peduncular cortex; DTT: dorsal tenia tecta; f: fornix; iLS: lateral septal nucleus, intermediate part; Ld: lambdoid septal zone; LS: lateral septum; MS: medial septum; NAc: nucleus accumbens; SFi: septofimbrial nucleus; SHi: septohippocampal nucleus; SHy: septohypothalamic nucleus; TS: triangular septal nucleus; VDB: vertical limb of the diagonal band of Broca; vLS: lateral septal nucleus, ventral part.

#### Tracing experiments

For retrograde tracing experiments, raphe images were acquired as z-stacks at 20× magnification on an inverted wide-field microscope (Leica DMi8), on sections sampled every 150 µm. Raphe neurons projecting to the vHP and/or LS were manually counted using the cell counter tool in FIJI. Tph2 immunostaining was used to define the anatomical architecture of the raphe nuclei, allowing for quantification of neuronal distribution across its subregions. The coordinates of detected neurons were extracted and their distribution along the antero-posterior, medio-lateral and dorso-ventral axes was analyzed using custom MATLAB scripts. For anterograde tracing experiment, images were acquired at 10× or 20× magnification. No quantitative analysis was performed on this dataset. For illustration purposes, all z-stacks images are shown as maximum projection.

### Statistics

Statistical tests were performed using OriginLab software. All bar graphs show mean ± SEM, box plots show median ± 1 SD. Mann-Whitney tests were used throughout the study and p values < 0.05 were considered statistically significant.

## Results

### VGLUT3^+^ axon terminal density, morphology and co-expression of 5-HT

We first characterized the distribution of VGLUT3 within the LS across its full rostro-caudal extent in WT mice, spanning from +1.70 mm to –0.46 mm relative to bregma. Using classic LS parcellations as an anatomical framework, VGLUT3^+^ terminals were observed throughout all major LS subdivisions—including the dorsal (dLS), intermediate (iLS), and ventral lateral septum (vLS), the medial septum/vertical limb of the diagonal band (MS/VDB), and the septofimbrial nucleus (SFi)—but their distribution displayed high regional heterogeneity (Figure 1) (Risold & Swanson, 1997; Swanson & Cowan, 1979).

Two main morphologies of VGLUT3^+^ terminals were consistently observed: PCBs (see Figure 1b), and individual axon terminals (or puncta). PCBs closely followed the soma contours of LS neurons, consistent with prior descriptions of VGLUT3- and monoamine-derived PCBs in limbic structures of mice, including the lateral septum (Riedel et al., 2008; Senft et al., 2021). Across LS subdivisions, PCBs were most abundant in the dLS, iLS, and SFi, where they formed dense, cell-encasing arrays. In the vLS, we observed a high density of individual VGLUT3*^+^* terminals mixed with PCBs. MS/DBB were dominated by individual puncta with fewer PCBs (Figure 1). At rostral levels (see bregma +1.70 to +0.98 mm in Figure 1A), VGLUT3 labeling formed a band-like PCB-rich zone, whereas at intermediate and caudal levels (see bregma +0.74 mm and beyond in Figure 1a), VGLUT3*^+^* PCBs became evenly distributed across LS sub-divisions. The caudal vLS displayed the highest density of VGLUT3 labeling, with individual puncta and PCBs forming a dense terminal field delineating the vLS border.

To obtain an unbiased overview of VGLUT3 organization across the lateral septal complex, we generated heatmaps of VGLUT3 puncta density and VGLUT3/5-HT colocalization across three bregma levels (Figure 2, bregma +1.42 mm, +0.38 mm and –0.22 mm). Colocalization with serotonergic markers was assessed because VGLUT3 and 5-HT co-transmission is a known feature of subsets of raphe-derived inputs, and because both neurotransmitters are present in pericellular baskets within septal circuits (Senft et al., 2021). Across all animals (n = 12, 6 males and 6 females), VGLUT3 density consistently appeared higher in the vLS compared to the dLS (see example heatmaps in Figure 2). Conversely, heatmaps of VGLUT3/5-HT colocalization suggested an increased density of VGLUT3^+^ axon terminal co-expressing 5-HT in the dLS, consistent with previous reports (Figure 2) (Senft et al., 2021; Shutoh et al., 2008). Heatmaps also highlighted a distinct profile of VGLUT3^+^ innervation of the septofimbrial nucleus (SFi, Figure 2, bregma –0.22 mm). Although the SFi exhibited only low to moderate VGLUT3^+^ terminal density, it consistently showed pronounced VGLUT3/5-HT colocalization, distinguishing it from neighboring LS territories.

**Figure 2.**
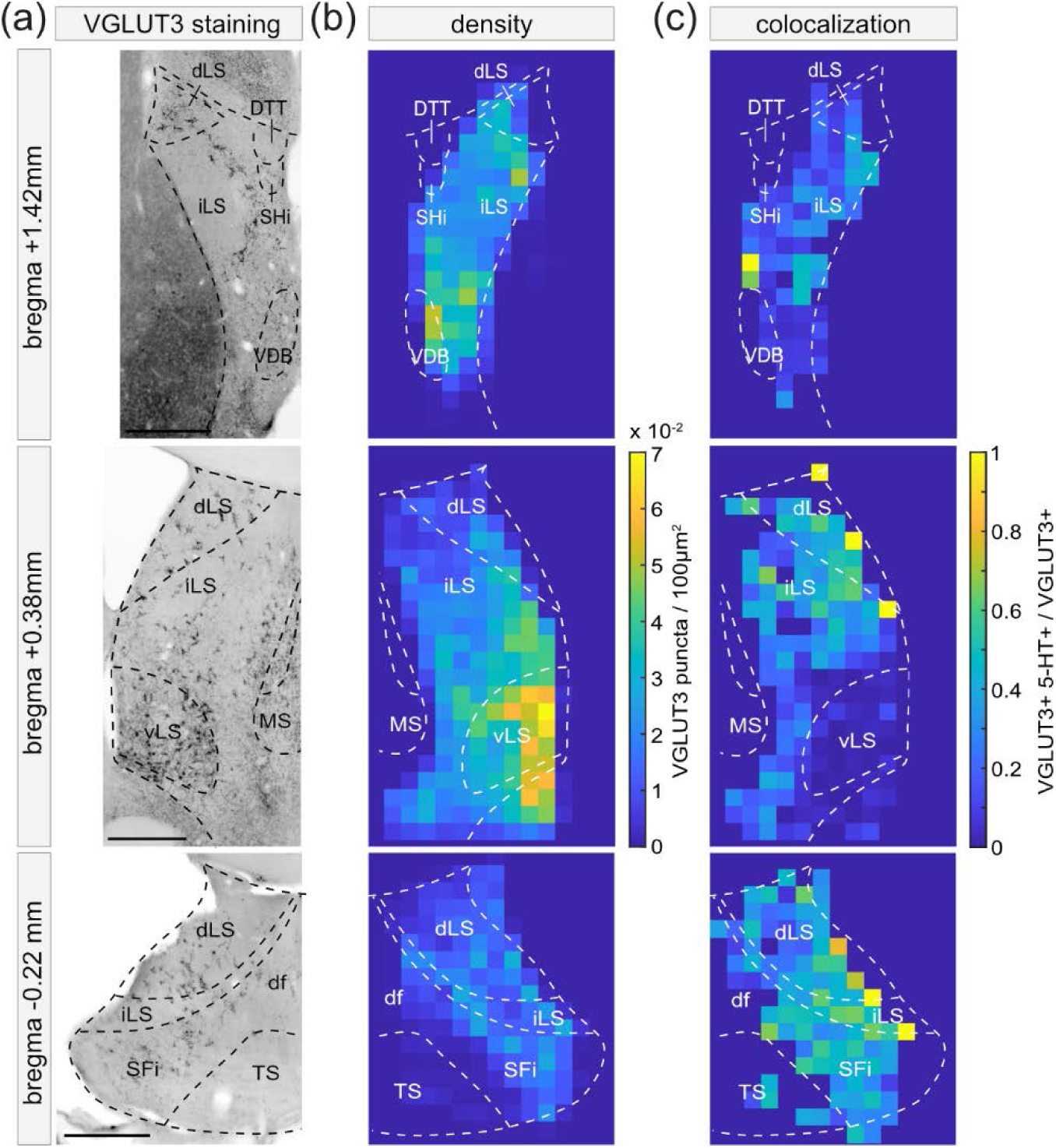
VGLUT3^+^ axon terminal density and colocalization with 5-HT in the lateral septum. (a) Immunohistochemical labeling of VGLUT3 in the LS of WT mice at three bregma levels (as indicated on the left). (b) Heatmaps showing the density of VGLUT3^+^ axon terminals in one example mouse at the same three LS levels. (c) Heatmaps showing colocalization with 5-HT in one example mouse at three LS levels. All scale bars 400 µm. n = 12 mice (6 female and 6 male).

Qualitative assessment of VGLUT3^+^ terminal density and colocalization with 5-HT at three representative levels suggests marked heterogeneity in the spatial distribution and neurochemical identity of VGLUT3^+^ inputs to the LS. We therefore performed quantitative analyses, focusing on a comparison between the dLS and vLS, in male and female mice (Figure 3). Averaged across LS rostro-caudal levels, the surface occupied by VGLUT3^+^ terminals was significantly higher in the vLS than in the dLS, with no differences between males and females (Figure 3a,b; males dLS: 5.24± 0.39%; females dLS: 5.09 ± 0.60%; males vLS: 15.62 ± 1.26%; females vLS: 18.66 ± 1.43%). When represented as a function of distance from bregma, VGLUT3 axon terminal density was highest the caudal half of the vLS, occupying up to 50% of the vLS surface. In contrast, in the dLS, density of VGLUT3 axon terminals remained relatively stable, occupying less than 20% of the dLS surface area. The rostro-caudal distribution of VGLUT3 density was similar between males and females (Figure 3c).

**Figure 3.**
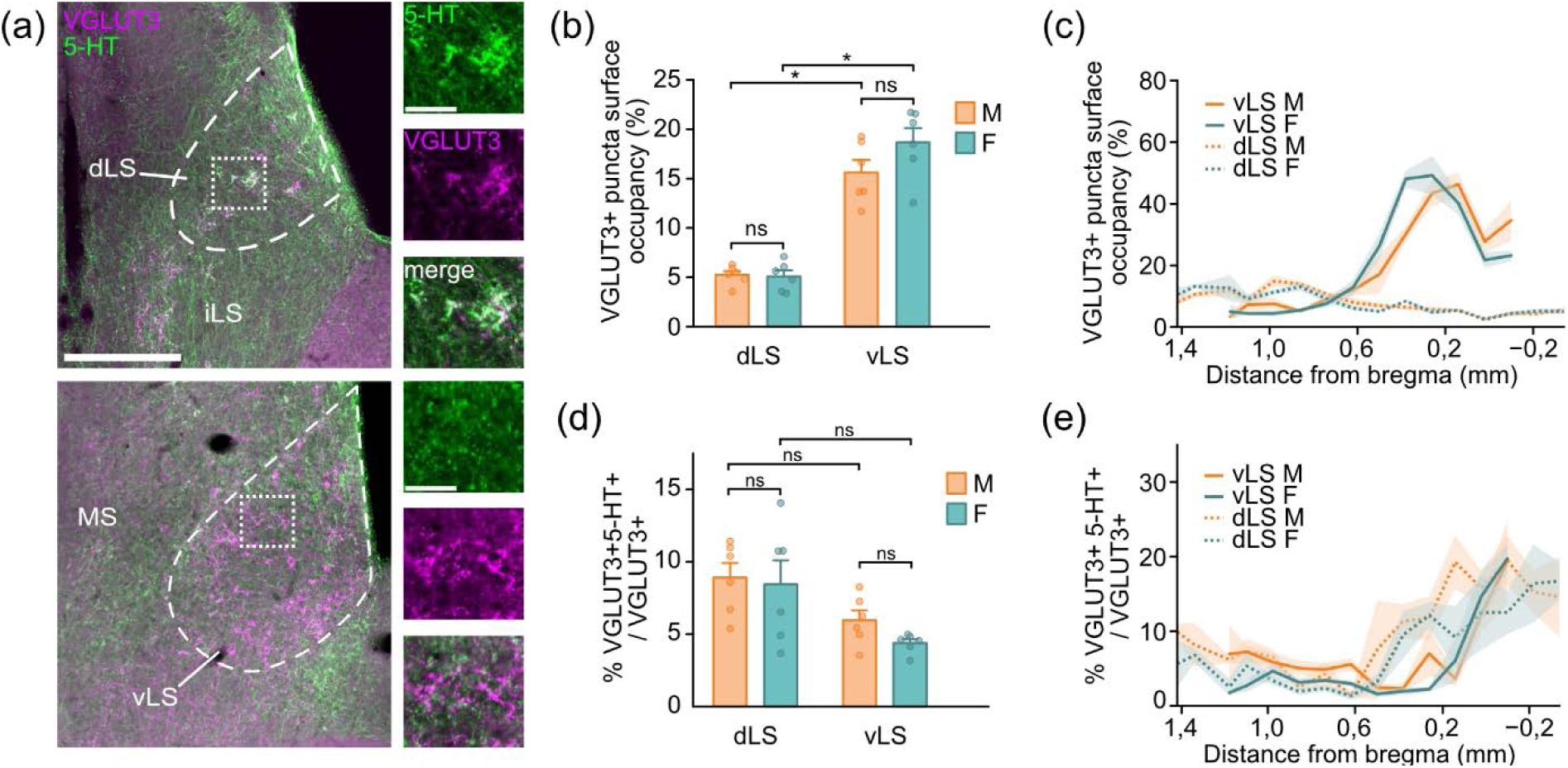
Distribution of VGLUT3 terminals in the dLS versus vLS and colocalization with 5-HT. (a) Immunohistochemistry for VGLUT3 (magenta) and 5-HT (green) in the dLS (top) and the vLS (bottom). Insets on the right show higher magnificiation overlays between VGLUT3 and 5-HT, illustrating the higher density of colocalization in the dLS compared to the vLS. (b) Density of VGLUT3 punctas in the dLS and vLS expressed as the proportion of vLS or dLS surface in male (n = 6) and female mice (n = 6). VGLUT3 axon terminal density was significantly higher in the vLS compared to dLS with no difference between sexes (males dLS vs vLS, *P* = 0.0051 ; females dLS vs vLS, *P* = 0.0051) (c) Same dataset as in b displayed as a function of the distance from bregma (in mm). (d) Proportion of VGLUT3 axon terminals colocalizing with 5-HT, over the total number of VGLUT3+ terminals in the dLS or vLS across all levels analyzed. (e) Distribution of VGLUT3 axon terminals colocalizing with 5-HT across the dLS and vLS rostro-caudal axis. Scale bar 200 µm (a), 20 µm (a - inset).

We next compared the proportion of VGLUT3^+^ terminals co-expressing 5-HT in the dLS and vLS of male and female mice. Averaged across all rostro-caudal levels, the proportion of VGLUT3+/5-HT+ boutons was higher in the dLS than in the vLS (Figure 3d, males dLS: 8.91 ± 1.001%; females dLS: 8.441 ± 1.65%; males vLS: 5.95 ± 0.69%; females vLS: 4.38 ± 0.27%). Analysis across the LS rostro-caudal axis further revealed that the proportion of colocalized axon terminals was highest in the caudal half of the LS for both the dLS and vLS (Figure 3e).

Altogether, VGLUT3^+^ inputs to the LS are spatially organized, displaying both a dorso-ventral gradient, with higher axon terminal density in the vLS relative to the dLS and SFi, and a rostro-caudal gradient, with the highest axon terminal density in the caudal half of the vLS. LS subdivisions are also contrasted in the neurochemical identity of VGLUT3 axon terminals, with the dLS and SFi containing a higher proportion of 5-HT co-expressing terminals whereas terminals in the vLS were predominantly non-serotonergic and therefore likely primarily glutamatergic.

### LS-projecting (VGLUT3^LS^) neurons are topographically organized within the raphe nuclei

To determine the topographical organization of raphe VGLUT3^+^ projections to the LS, we injected a Cre-dependent retrograde AAV into the LS of VGLUT3-Cre mice and examined the distribution of retrogradely labeled neurons throughout raphe sub-regions (Figure 4a). Viral spread encompassed a large portion of the LS while largely sparing the medial septum (MS) and vertical limb of the diagonal band (MS/VDB; Figure 4b), ensuring that retrograde labeling remained specific to LS-projecting populations.

**Figure 4.**
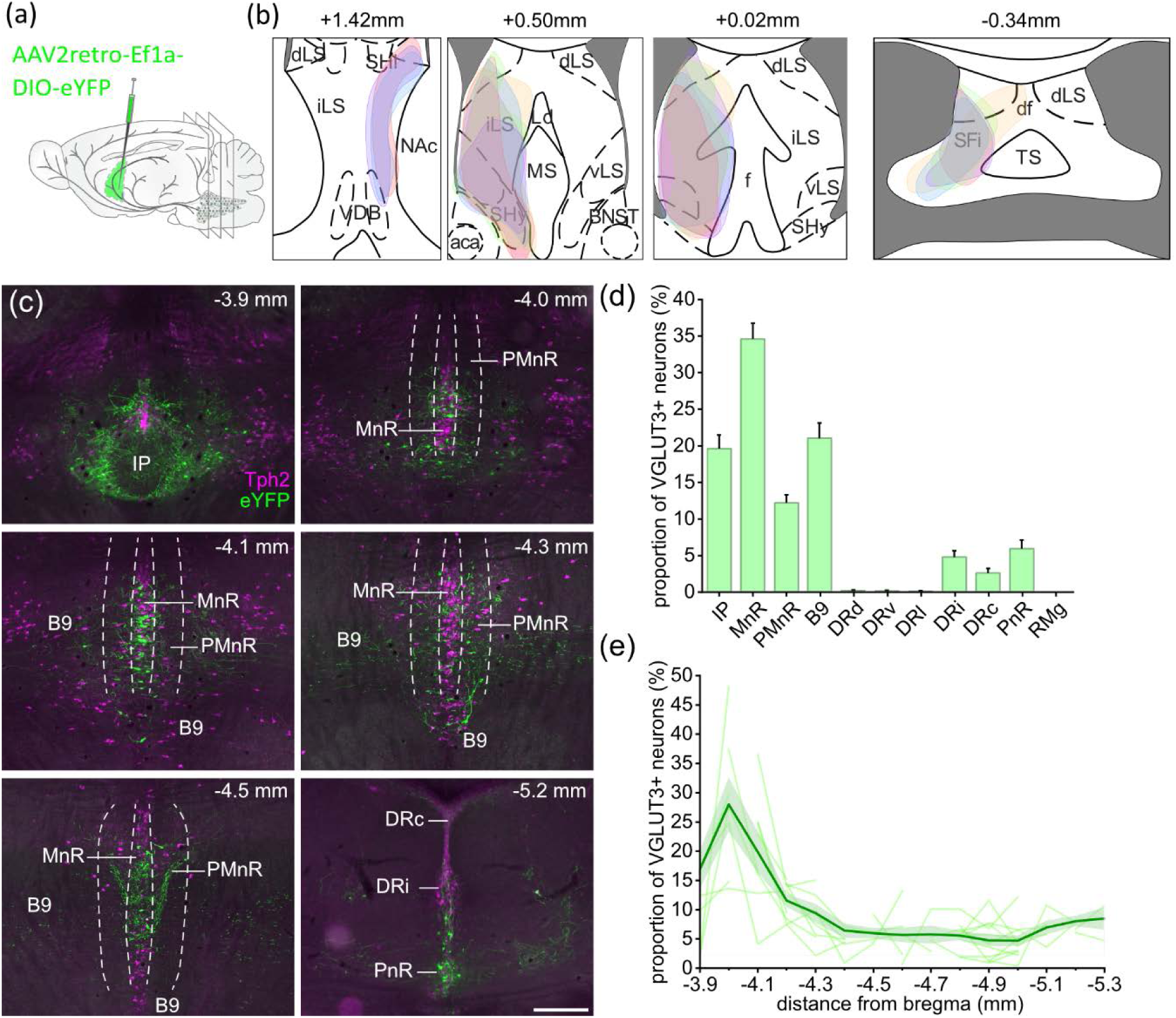
Distribution of LS-projecting VGLUT3^+^ neurons within the raphe nuclei. (a) Schematic of Cre-dependent viral vector injection in the LS of VGLUT3-Cre mice. (b) Schematic representation of LS areas where eYFP expression was detected in axon terminals and PCBs, shown for n = 6 mice. (c) Retrograde expression of eYFP (green) and localization of VGLUT3^LS^ neurons in different raphe sub-regions, delineated using Tph2 expression (magenta). (d) Distribution of VGLUT3^+^ neurons in raphe subregions. The IP was included in the quantification due to the high density of VGLUT3^LS^ neurons. Results are expressed as the proportion of the total number of VGLUT3 neurons across all raphe sub-regions. (e) Rostro-caudal distribution of VGLUT3^+^ neurons, shown as a function of distance from bregma (in mm). Scale bar: 300µm. n = 12 mice (7 males, 5 females). aca: anterior commissure, anterior part; B9: supralemniscal nucleus; BNST: bed nucleus of stria terminalis; df: dorsal fornix; dLS: lateral septum, dorsal part; DP: dorsal peduncular cortex; DRc: caudal part of the dorsal raphe; DRi: interfascicular part of the dorsal raphe; DRl: lateral part of the raphe; DRv: ventral part of the dorsal raphe; DTT: dorsal tenia tecta; f: fornix; iLS: lateral septum, intermediate part; IP: interpeduncular nucleus; Ld: lambdoid septal zone; MnR: median raphe; MS: medial septum; PMnR: paramedian raphe; PnR: pontine raphe nucleus; Rmg: raphe magnus; SFi: septofimbrial nucleus; SHi: septohippocampal nucleus; TS: triangular septal nucleus; VDB: nucleus of the vertical limb of the diagonal band; vLS: lateral septum, ventral part.

Across all rostro-caudal levels of the raphe nuclei, retrograde GFP labeling revealed that LS-projecting VGLUT3^+^ neurons were predominantly localized within the median raphe region (MRR, corresponding to median raphe [MnR] and paramedian raphe [PMnR]) and the supralemniscal B9 cell group (Figure 4 c,d). More caudal raphe subregions, including DRi, DRc, and the pontine raphe (PnR), contributed to smaller fractions of LS-projecting neurons (Figure 4d). Additional LS-projecting neurons were detected within the lateral subdivision of the IP (Figure 4 c,d), a region closely related to the MRR and connected with raphe neuromodulatory circuits (Kawai et al., 2022; Lima et al., 2017). Along the rostro-caudal axis, VGLUT3^LS^ neuron distribution revealed a marked enrichment in the most rostral part of the raphe nuclei, corresponding to the MnR, PMnR, and B9 cell group (Figure 4e).

Closer inspection of the most rostral levels of the raphe (bregma −4.0 to −4.1 mm) revealed an unexpected cluster of VGLUT3^+^ somata located ventral to the MRR (Figure 4c and Figure 5a). To determine whether this ventral cluster was located in the caudal part of the IP, we performed choline acetyltransferase (ChAT) immunostaining. Dense cholinergic innervation from the medial habenula sharply delineates the dorsal and intermediate sub-regions of the IP (Frahm et al., 2015; Hsu et al., 2013). At levels corresponding to bregma -4.0 mm and beyond, ChAT immunostaining was almost completely absent, suggesting that this subset of VGLUT3^LS^ neurons does not belong to the IP (Figure 5a).

**Figure 5.**
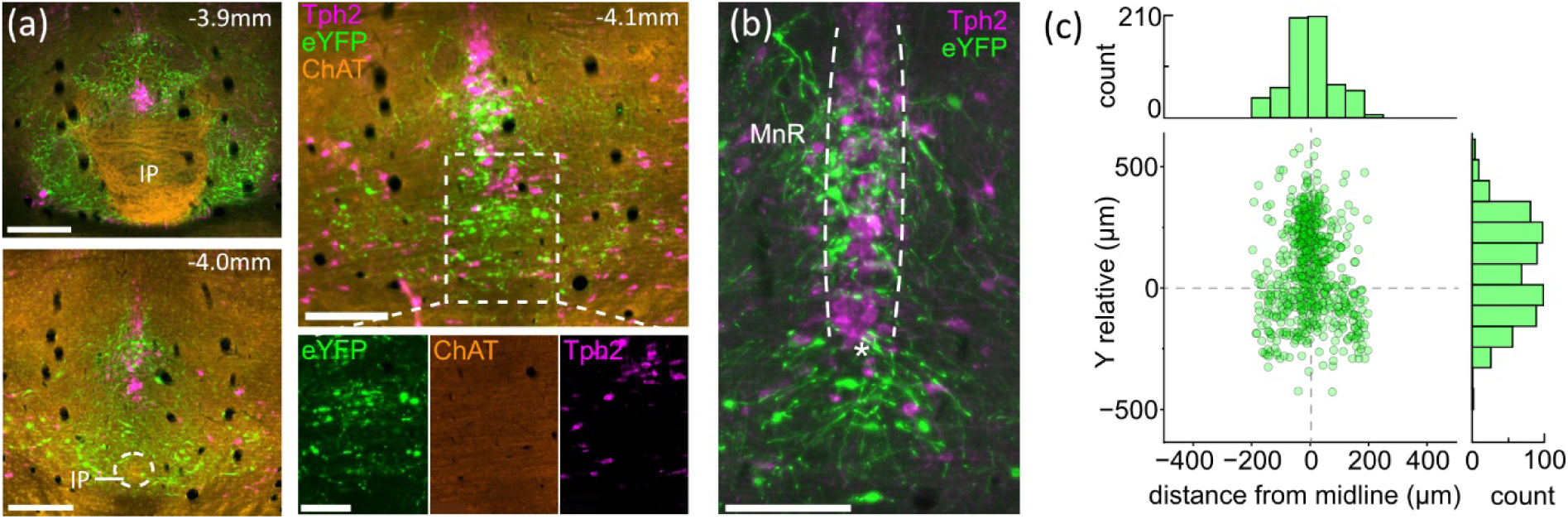
LS-projecting neurons distribution unveils a VGLUT3+/Tph2-negative neuronal population ventral to the MnR. (a) Immunohistochemistry against Tph2 and Choline Acetyltransferase (ChAT) delineating the boundaries of the MRR and the IP. The inset shows a higher-magnification imageof VGLUT3^LS^ neurons found ventral to the MRR within a ChAT-negative region. (b) Example image showing the MnR, delineated by the dotted lines, the reference point that was chosen to identify the ventral boundary the MRR (star) and eYFP-positive VGLUT3^LS^ neurons located within and ventral to the MRR. (c) Spatial distribution of VGLUT3^LS^ neurons relative to the ventral boundary of the MRR. Bregma levels -4.0, -4.1, -4.2 mm were included in the analysis. Scale bar: 200 µm (a [left, top and bottom], b), 100 µm (a [right]- inset bottom panel). n = 11 mice (6 males, 5 females).

The spatial distribution of VGLUT3^LS^ neurons within the MRR was analyzed by quantifying the position of VGLUT3^LS^ neurons relative to the ventral boundary of the MRR, defined here as the position of the most ventral Tph2-positive neuron of the dense neuron population of the MRR (Figure 5 b,c). As expected, a subset of VGLUT3^LS^ neurons were found along the midline and within the MRR. However, a distinct population was positioned ventral to MRR Tph2+ neurons and displayed a more dispersed medio-lateral distribution. The dorso–ventral segregation was consistent across mice (n = 11), suggesting that the ventral MRR population is a reproducible anatomical feature of the LS-projecting VGLUT3 system. Overall, these findings indicate the existence of a VGLUT3^+^ subpopulation of LS-projecting neurons, ventral to the MRR, never described in the literature before (Okaty et al., 2015; Ren et al., 2019; Senft et al., 2021; Sos et al., 2017).

Together, these results demonstrate that VGLUT3^LS^ neurons form a spatially organized system within the raphe, characterized by: (1) a strong rostral bias centered in the MRR/B9 regions; (2) a dorso–ventral specific organization within the MRR; and (3) a distinct ventral MRR population that is anatomically and neurochemically separable from both the dorsally located MRR cluster and the IP. This organization mirrors the developmental and molecular heterogeneity of the raphe nuclei and suggests that the LS receives convergent inputs from several distinct subsets of median raphe VGLUT3 neurons.

### VGLUT3^LS^ and VGLUT3^vHP^ neurons are spatially segregated within the raphe nuclei

The LS shares functional roles with the ventral hippocampus (vHP) in feeding regulation and affective behaviors (Albert & Wong, 1978; Blanchard et al., 1979; Carus-Cadavieco et al., 2017; Kjelstrup et al., 2002; Kosugi et al., 2021; Sweeney & Yang, 2015). We have previously characterized VGLUT3+ raphe inputs to the vHP (Fortin-Houde et al., 2023), raising the possibility that vHP-projecting VGLUT3 neurons (VGLUT3^vHP^) and VGLUT3^LS^ neurons partially overlap, with subsets of VGLUT3^+^ raphe neurons sending collateral projections to both LS and vHP. To compare the organization of VGLUT3^vHP^ and VGLUT3^LS^ populations, we performed dual retrograde viral tracing by injecting Cre-dependent retrograde viral vectors into both structures (Figure 6a).

**Figure 6.**
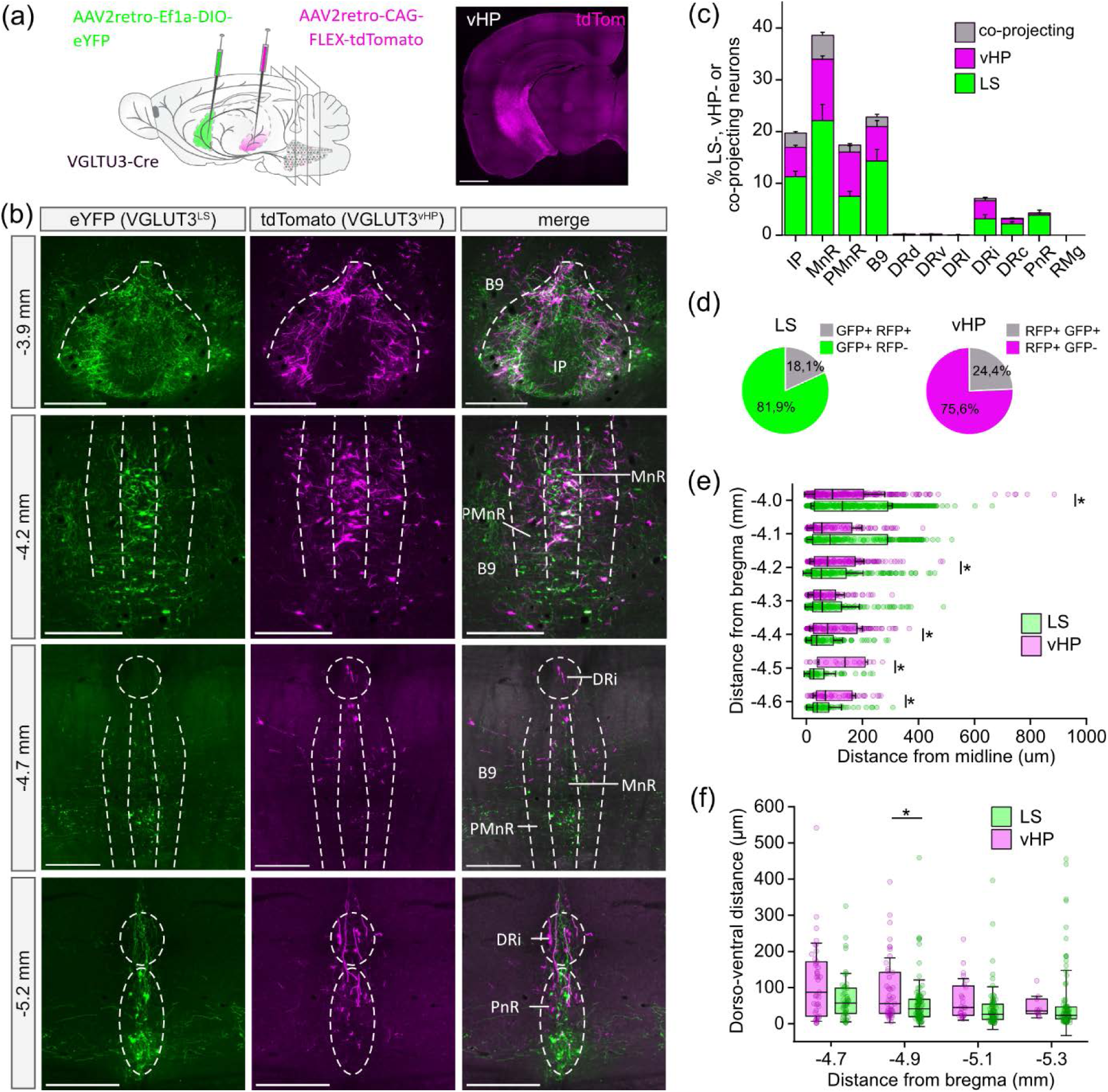
VGLUT3^+^ neurons projecting to LS and vHP are spatially segregated within the raphe nuclei, with limited overlap between the two populations. (a) Injection schematic of cre-dependent viral vectors in the LS and vHP of VGLUT3-cre mice. Image on the right shows an example of the vHP expression in injection site. (b) Immunohistochemical labeling of eYFP (green) and tdTomato (magenta), identifying VGLUT3^LS^ and VGLUT3^vHP^ neurons respectively. Images show their rostro-caudal and dorso-ventral distribution at different raphe levels (distance from bregma indicated on the left). (c) Proportion of VGLUT3^+^ neurons projecting to the LS (green), vHP (magenta) or both (grey) per region (raphe subregions and IP) out of all LS- and vHP-projecting VGLUT3 neurons. (d) Pie charts depicting the proportion of co-projecting neurons within VGLUT3^LS^ (left) or VGLUT3^vHP^ (right) populations. (e) Medio-lateral distribution of LS- and vHP-projecting neurons in the MRR, at different distances from bregma. Each point represents a VGLUT3^+^ soma (LS-projecting or vHP-projecting only, co-projecting neurons were not represented for this quantification). For all box-plots, the box represents 25-75 percentile with error bar of 1 SD and line represents the median. Bregma –4.0 mm: *P* = 0.0399; bregma –4.2 mm: *P* = 0.0251; bregma –4.4 mm: *P* = 0.0030; bregma –4.5 mm: *P* = 0,0005; bregma –4.6 mm: *P* = 0,0030. (f) Dorso-ventral distribution of LS- and vHP-projecting neurons in the MRR at different distances from bregma. Distance along the midline was calculated for each VGLUT3^+^ soma (LS-projecting or vHP-projecting only). Bregma -4.9 mm: *P* = 0,0169. Scale bar: 1000 µm (a), 300 µm (b). n = 12 mice (7 males, 4 females, 1 WT control).

We first examined the distribution of VGLUT3^vHP^ and VGLUT3^LS^ neurons within the raphe sub-regions. Immunostaining against eYFP and tdTomato, identifying VGLUT3^LS^ and VGLUT3^vHP^ neurons respectively, revealed that the two populations were spatially segregated (Figure 6b,c). Although VGLUT3^vHP^ and VGLUT3^LS^ neurons were predominantly found within the same four raphe sub-regions (IP, MnR, PMnR and B9 group), they displayed limited overlap within these sub-regions (Figure 6c), consistent with a largely divergent and neurochemically segregated organization of projection patterns, rather than strictly parallel, non-collateralized streams (Senft et al., 2021). Quantification of the total proportion of co-projecting neurons across all raphe subregions showed that 24.4% of vHP-projecting neurons co-projected to the LS, and conversely 18.1% of LS-projecting neurons also projected to the vHP (Figure 6d). These numbers point to the existence of a shared VGLUT3^+^ subsystem within the raphe nuclei and may provide an anatomical basis for coordinated regulation of LS and vHP functions mediated by the raphe nuclei.

We also interrogated whether VGLUT3^vHP^ and VGLUT3^LS^ neurons displayed distinct spatial distributions. Quantification of medio-lateral distribution demonstrated that at rostral levels of the raphe nuclei (bregma -4.0 mm), LS-projecting neurons were located significantly more laterally than vHP-projecting neurons. This pattern reversed at more caudal levels (bregma -4.4 to -4.6 mm), with LS-projecting neurons shifting closer to the midline and vHP-projecting neurons occupying comparatively lateral positions (Figure 6e). At the most caudal levels of the raphe (bregma -4.7 to -5.3 mm), the two populations also displayed dorsoventral segregation, with VGLUT3^vHP^ neurons occupying significantly more dorsal positions, while VGLUT3^LS^ neurons were enriched ventrally (Figure 6f). This differential distribution is consistent with earlier reports demonstrating that VGLUT3^+^ neurons in the MnR and PMnR form longitudinally segregated channels targeting discrete forebrain structures, including the septum, hippocampus and cortex (Fortin-Houde et al., 2023; Senft et al., 2021).

To further investigate the neurochemical identity of these two projection pathways, we quantified serotonergic marker expression in VGLUT3^LS^ and VGLUT3^vHP^ neurons (Figure 7). Part of VGLUT3 raphe neurons share their developmental lineage with 5-HT neurons, under control of the Pet1 transcription factor. Pet1-positive neurons comprise neurochemically heterogenous populations, including predominantly 5-HT neurons expressing high levels of the 5-HT biosynthesis enzyme Tph2 and low levels of VGLUT3, predominantly glutamatergic neurons characterized by VGLUT3-high/Tph2-low, and neurons co-expressing VGLUT3 and Tph2 (Alonso et al., 2013; Okaty et al., 2015; Pelosi et al., 2014; Ren et al., 2019). Recent studies have identified that part of LS-projecting raphe neurons display a mixed VGLUT3/5-HT phenotype (Henderson et al., 2024; Senft et al., 2021). We therefore quantified the proportion of identified VGLUT3^LS^ neurons co-expressing the serotonergic marker Tph2. Immunohistochemistry revealed a marked difference in serotonergic identity: along the rostro-caudal axis, VGLUT3^LS^ neurons display minimal Tph2 co-expression, whereas VGLUT3^vHP^ neurons show a substantially higher proportion of Tph2^+^ somas (Figure 7 a,b). Across all raphe sub-regions, we found significantly higher proportions of Tph2 co-expression in VGLUT3^vHP^ neurons compared to VGLUT3^LS^ neurons, confirming a robust dissociation in the neurochemical phenotype between these two pathways (Figure 7c; VGLUT3^vHP^: 27.9 ± 3.0%; VGLUT3^LS^: 5.3 ± 0.9%).

**Figure 7.**
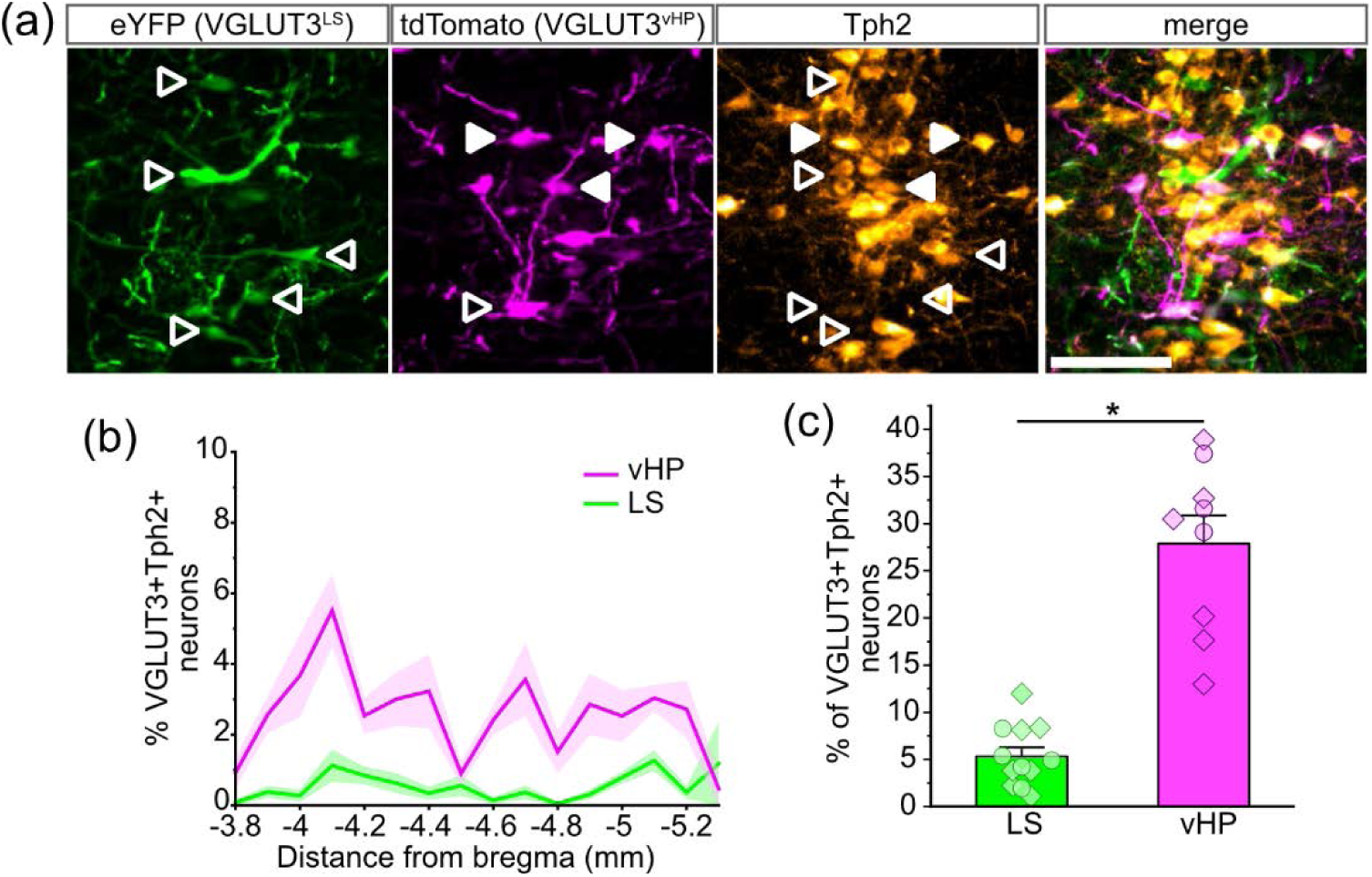
VGLUT3^LS^ and VGLUT3^vHP^ neuron co-expression with Tph2. (a) Immunohistochemistry against eYFP, tdTomato and Tph2 in the MRR. Empty arrowheads indicate VGLUT3^LS^ or VGLUT3^vHP^ neurons that do not co-express Tph2, filled arrowheads indicate somas co-expressing Tph2. (b) Proportion of Tph2-positive VGLUT3^vHP^ (magenta) and VGLUT3^LS^ (green) neurons along the rostro-caudal axis of the raphe nuclei. (c) Total proportion of Tph2 co-expression within the raphe nuclei, calculated as a fraction of all VGLUT3^LS^ (green) or VGLUT3^vHP^ (magenta). Male mice are indicated with diamonds, females with circles. vHP: n = 9 mice (6 males, 3 females), LS: n = 12 (7 males, 5 females), *P* = 0.0081. Scale bar in (a): 100 µm.

Together, these results show that VGLUT3^+^ neurons projecting to the LS and vHP form largely independent subcircuits within the raphe nuclei. Their contrasting medio-lateral and dorso-ventral distributions highlight a refined topographical organization, while the presence of co-projecting neurons identifies a subset of VGLUT3^+^ population capable of influencing both structures. The pronounced difference in Tph2 co-expression indicates that these pathways are not only anatomically but also neurochemically distinct, support the idea that Pet1-derived raphe neurons form multiple parallel, functionally differentiated systems.

### VGLUT3^+^ afferents originate in various brain regions that tile distinct LS domains

Our retrograde tracing experiments revealed that VGLUT3^+^ axon terminals in the LS do not originate solely from the raphe nuclei (Figure 8). Part of VGLUT3 inputs originated from VGLUT3^+^ neurons found in the BNST, where retrogradely labeled eYFP-expressing neurons were densely present (Figure 8 b,c). This result is in agreement with literature demonstrating a strong BNST to LS projection that contributes to stress-related, autonomic, and motivational functions (Dong & Swanson, 2004; Maita et al., 2021). A second prominent input originated from the NI and the adjacent PCG, with neurons expressing the neuropeptide relaxin-3 delineating the NI (Figure 8 d-f). This pattern is consistent with previous reports showing that NI neurons, particularly relaxin-3 and glutamatergic subpopulations, target the LS and may regulate arousal, stress integration, and hippocampal theta modulation (Goto et al., 2001; Wang & Morales, 2009). Together, the BNST, MRR/IP and NI/PCG form the three major sources of VGLUT3^+^ inputs to the LS.

**Figure 8.**
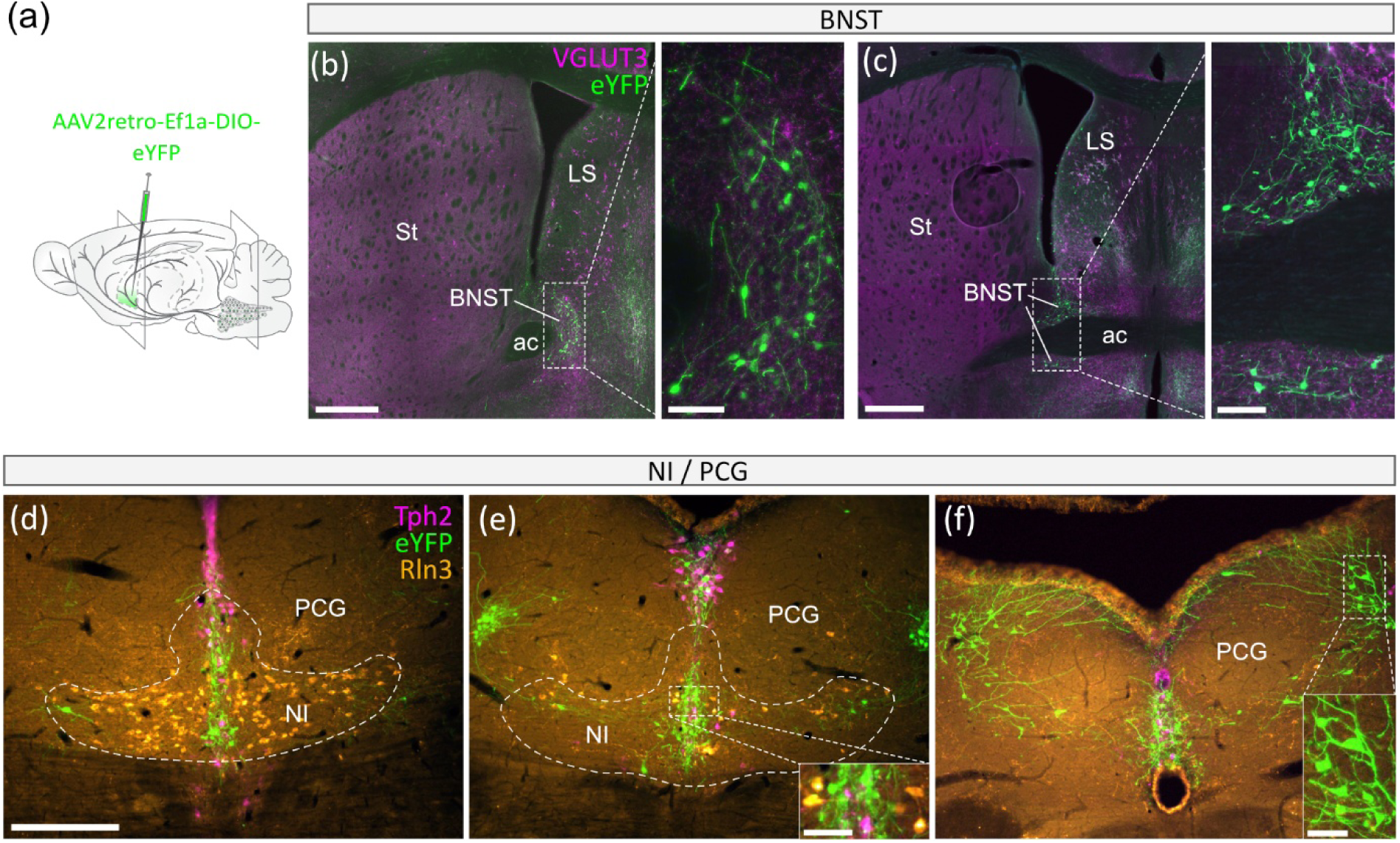
Retrograde tracing from LS reveals multiple brain regions contributing to VGLUT3^+^ inputs to LS. (a) Schematic illustrating injection of a retrograde Cre-dependent AAV in the LS of VGLUT3-Cre mice. (b) eYFP expression in retrogradely labelled VGLUT3 neurons (green) and VGLUT3 immunoreactivity (magenta) in the BNST. (c) Same labeling shown at a more caudal level of the BNST. (d-f) Immunohistochemistry detecting eYFP (green), Tph2 (magenta) and Relaxin-3 (Rln3, orange) in the NI/PCG. Scale bars: 500 µm (b and c), 100 µm (insets in b and c), 300 µm (d, e, f), 50 µm (insets in e and f). ac: anterior commissure; BNST: bed nucleus of stria terminalis; LS: lateral septum; NI: nucleus incertus; PCG: pontine central gray; St: striatum.

We next examined, using anterograde tracing, how VGLUT3^+^ inputs from the BNST, MnR/IP and NI/PCG distribute in the LS (Figure 9). Injections targeting NI and PCG were pooled because even low volume injections did not allow selective labeling of either region alone. Following injection in the NI/PCG, we observed abundant axon fiber and terminals throughout the rostro-caudal axis of the LS (Figure 9a). Most consisted of thin axons with multiple varicosities that colocalized with VGLUT3 immunostaining. The highest density of VGLUT3^+^ NI/PCG inputs was found in the rostral dLS (see bregma +1.42 mm) and throughout the vLS at all levels examined. To a lesser extent, we also observed isolated axon terminals and thicker, smooth fibers throughout the LS. We observed virtually no PCBs arising from NI/PCG inputs to the LS. These findings are in line with previous anatomical studies showing that NI neurons, including relaxin-3-containing populations, project to the septal region and hippocampal formation via the medial forebrain bundle (Goto et al., 2001; Ryan et al., 2011).

**Figure 9.**
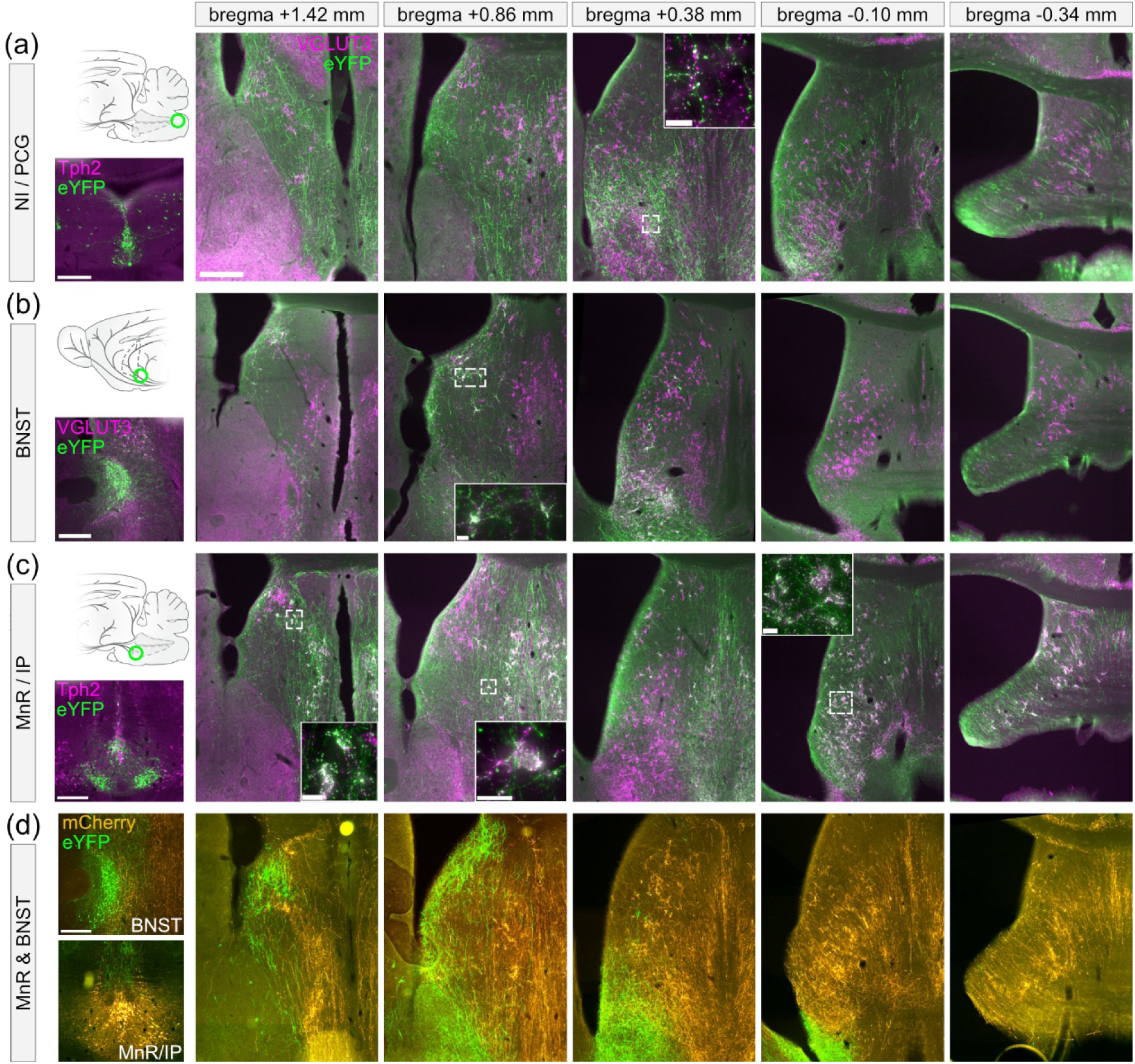
BNST, MnR/IP and NI/PCG VGLUT3^+^ inputs tile distinct LS regions. (a) Left: schematic illustrating injection site of anterograde Cre-dependent AAV in VGLUT3-cre mice, driving the expression of eYFP in the NI/PCG (top), and image of the injection site (bottom). Tph2 staining is in magenta. Right: example images showing the extent of VGLUT3^+^ NI/PCG innervation of the LS at various levels, indicated on the top as the distance from bregma. (b) Same for BNST injections. VGLUT3 staining is in magenta. (c) Same for MnR/IP injections. Tph2 staining is in magenta. (d) Left: images of injections sites following two-color anterograde tracing of BNST (top) and MnR/IP (bottom) VGLUT3^+^ inputs. Right: anterograde expression of eYFP and mCherry in VGLTU3^+^ terminals at different bregma levels of the LS. For all injections, consistency of the innervation pattern was verified in minimally n = 3 mice. Scale bar: 300 µm (a-d), 20 µm (insets).

Compared to NI/PCG, injections in the BNST resulted in a more modest density of axon terminals in the LS. At rostral levels, BNST inputs formed VGLUT3^+^ PCBs that were confined to the latero-dorsal LS, while largely avoiding the MS and midline regions (Figure 9b). BNST-originating PCBs were restricted to the rostral half of the dLS, with no BNST inputs detected in this area beyond bregma +0.38 mm. BNST-originating PCBs also densely innervated the vLS at intermediate levels on the rostro-caudal axis (see bregma +0.38 mm). This pattern is consistent with a classical tracing study showing that posterior BNST sends dense, topographically organized projections to rostral lateral septal regions, particularly in dorso-lateral and ventro-lateral subdivisions (Dong & Swanson, 2004).

In contrast, injections targeting the MnR/IP resulted in dense and widespread coverage across the entire rostro-caudal extent of the LS (Figure 9c). Terminals from MnR/IP formed numerous VGLUT3^+^ PCBs, with a particularly high density in the iLS and vLS. At rostral levels (bregma +1.42 mm and +0.86 mm), afferents appeared to mostly avoid the dLS and rather concentrated close to midline and flanking the medial septum. At caudal levels (bregma +0.38 mm to -0.34 mm), MnR/IP VGLUT3^+^ inputs virtually represented the entirety of VGLUT3 axon terminals. When considered alongside the BNST innervation pattern, our results showed striking complementarity of BNST- and MnR/IP-derived VGLUT3 inputs, with BNST accounting for PCBs in the rostral parts of the dLS and vLS, while MnR/IP projections covered the iLS across the entire rostro-caudal extent, and ventral and lateral divisions of the caudal LS. This organization is consistent with previous work showing that MnR VGLUT3^+^ neurons heavily innervate the LS and MS (Fortin-Houde et al., 2023). This complementarity distribution of BNST vs MnR/IP inputs to the LS was further visualized using double anterograde tracing from the BNST and MnR/IP, illustrating preferential targeting of the latero-dorsal rostral LS by BNST projections, whereas MnR/IP axons formed PCBs in all remaining LS subdivisions (Figure 9d).

Together, these results demonstrate that the LS is tiled by VGLUT3^+^ inputs from at least three anatomically and morphologically distinct sources. Each source targeting partially overlapping but topographically biased LS domains with characteristic terminal phenotypes (Figure 10),

**Figure 10.**
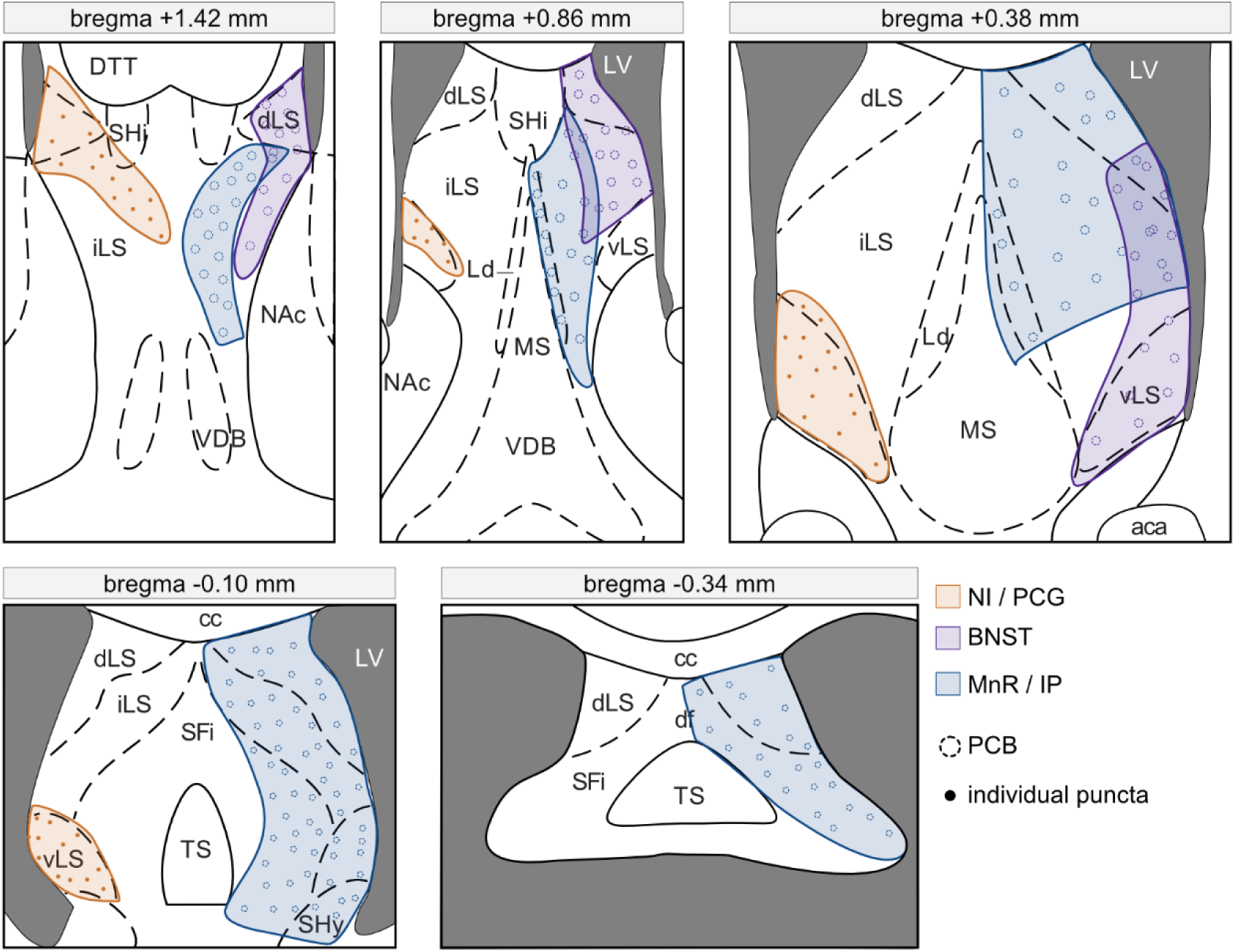
Afferents from various brain regions tile the LS. Schematic summarizing the topography of VGLUT3^+^ inputs to LS along its rostro-caudal axis. Inputs arising from BNST (purple), NI/PCG (orange) and MnR/IP (blue) are shown, alongside the morphology of axon terminals, shown as circle (PCBs) or dots (individual punctas). Projections were illustrated unilaterally for clarity purposes. aca: anterior commissure, anterior part; cc: corpus callosum; D3V: third ventricule, dorsal part; df: dorsal fornix; dLS: lateral septal nucleus, dorsal part; DTT: dorsal tenia tecta; iLS: lateral septal nucleus, intermediate part; ; Ld: lambdoid septal zone; LV: lateral ventricules; MS: medial septum; NAc: nucleus accumbens; SFi: septofimbrial nucleus; SHi: septohippocampal nucleus; SHy: septohypothalamic nucleus; TS: triangular septal nucleus; VDB: vertical limb nucleus of the diagonal band of Broca; vLS: lateral septal nucleus, ventral part.

## Discussion

This study provides the first comprehensive anatomical characterization of VGLUT3^+^ inputs to the LS and their source populations within the raphe nuclei and other limbic and brainstem regions. Three main findings emerge: (1) VGLUT3^+^ terminals exhibit a pronounced dorso-ventral organization within the LS, with substantially higher density in the ventral subdivision and higher 5-HT colocalization in the dLS; (2) VGLUT3-expressing neurons projecting to the LS and those projecting to the vHP are spatially segregated within the raphe nuclei, with limited but noteworthy overlap; and (3) the LS receives VGLUT3^+^ afferents from multiple sources, including the raphe nuclei, the BNST and NI/PCG, each displaying distinct terminal organization and morphology. Together, these results refine the anatomical framework of VGLUT3 inputs to the LS and reveal organizational principles that were previously unresolved.

The marked dorso-ventral heterogeneity in VGLUT3^+^ puncta density and colocalization with 5-HT confirms and extends recent findings, describing higher colocalization between VGLUT3 and 5-HT in the dLS (Senft et al., 2021). Part of VGLUT3/5-HT co-expressing axon terminals in the dLS have been shown to arise from Pet1-derived raphe neurons from rhombomere 2. Yet, r2-derived Pet1 neurons do not account for all VGLUT3^+^ terminals in the LS, suggesting that neurons from other lineages also innervate this structure (Senft et al., 2021). Our results confirm these findings, showing that although a majority of VGLUT3^LS^ neurons were concentrated in the MRR, high proportion of neurons can also be found in the B9, IP and to a lesser extent in caudal structures of the raphe nuclei (DRi, DRc and PnR,) that altogether contain Pet1-derived neurons of diverse rhombomeric origins.

The distinct distribution and neurochemical identity of VGLUT3^+^ terminals across the dLS and vLS mirror the established anatomical and functional organization of LS subdivisons. Consistent with this segregation, the dLS has been primarily associated with spatial coding, navigation, avoidance behaviors, and conditioned fear, whereas the vLS is more strongly implicated in motivational, emotional, social, and anxiety-related processes (Patel, 2022; Sheehan et al., 2004; van der Veldt et al., 2021; Wirtshafter & Wilson, 2021). Recent studies have further emphasized the role of the vLS in anxiety and affective regulation (Liu et al., 2024; Peng et al., 2024; Rashid et al., 2025; Yeates et al., 2022). Our findings also extend classical anatomical studies demonstrating that LS functional compartmentalization is supported by organized afferent connectivity, with brainstem inputs, including raphe-derived fibers, preferentially innervating ventral regions, while hippocampal and hypothalamic afferents more strongly target dorsal subdivisions (Risold & Swanson, 1997; Swanson & Cowan, 1979). Together, these observations suggest that VGLUT3 expression and serotonergic colocalization are integrated into the broader dorso-ventral specialization of LS circuits.

Many VGLUT3^+^ terminals are organized in PCBs with differences between subdivisions of the LS, the dLS and iLS having higher proportions of PCBs relative to the vLS. VGLUT3^+^ PCBs were initially reported in the rat LS, and subsequent studies identified the MnR as the main source of LS PCBs in mice (Herzog et al., 2004; Riedel et al., 2013; Riedel et al., 2008; Senft et al., 2021). Our results extend these findings by highlighting species differences in the organization of VGLUT3^+^ PCBs within the LS. In rats, VGLUT3-immunoreactive PCBs display a highly ordered, laminar and band-like arrangement that spans the LS in oblique stripes, largely independent of canonical septal divisions (Riedel et al., 2008). These PCBs are most prominent in dorsal and intermediate subdivisions, whereas vLS and SFi lack well-defined PCBs and instead contain a diffuse VGLUT3^+^ fiber arrangement (Herzog et al., 2004; A. Riedel et al., 2008; A. Riedel et al., 2013). In contrast, our results in mice reveal a less stereotyped but regionally biased organization in which PCBs are preferentially enriched in the dLS and iLS but are also detectable in the SFi and to a lesser extent, in the vLS, coexisting with abundant isolated VGLUT3^+^ terminals. Moreover, whereas rat studies suggest predominantly brainstem origins for LS-targeting VGLUT3^+^ inputs, our results in mice demonstrate that VGLUT3^LS^ neurons arise from a broader diversity of brain regions.

We found that the largest source of VGLUT3^+^ inputs to the LS originates from the IP, MnR and B9, together contributing to a widespread pattern of VGLUT3^+^ innervation across all LS subdivisions. Although not part of the raphe nuclei proper, the IP displays strong anatomical connectivity with the MnR and an established role in modulating septo-hippocampal pathways (Frahm et al., 2015; Hsu et al., 2013). The vast majority of IP/MnR/B9-derived inputs formed PCBs within the LS. In the MS, we observed axons with VGLUT3^+^ varicosities but no PCBS, consistent with previous observations (Fortin-Houde et al., 2023). Raphe/IP-derived inputs largely avoided the most rostral levels of the dLS, which instead contained VGLUT3^+^ PCBs originating from the BNST. In contrast, BNST-derived inputs formed fewer PCBs, and preferentially targeted lateral and dorsal LS subdivisions, with more limited innervation of the vLS. This pattern of connectivity supports the idea that the BNST-LS system forms a major component of extended amygdala circuits involved in stress, anxiety and defensive behaviors (Hong - Wei Dong & Larry W Swanson, 2004). Together, VGLUT3^+^ inputs from the raphe/IP and BNST accounted for virtually all PCBs in the LS and showed striking complementarity in their spatial organization. A third source of VGLUT3^+^ inputs to the LS arose from the NI and PCG. These regions, implicated in arousal, attention, valence and anxiety regulation contain a small and poorly characterized population of VGLUT3^+^ neurons (Goto et al., 2001; Ryan et al., 2011; Xiao et al., 2023). NI/PCG VGLUT3^+^ inputs were abundant, covering the rostral dLS and mid- to caudal vLS, and appeared almost exclusively as individual puncta or axon varicosities rather than PCBs. Together, these findings reveal a complex anatomical organization of VGLUT3^+^ inputs within the LS and outline a rich and multimodal glutamatergic network.

Our study further reveals the spatial segregation between MRR VGLUT3+ neurons projecting to the LS and those projecting to the vHP, consistent with previous reports showing that MRR outputs to limbic structures are topographically organized (Fortin-Houde et al., 2023; Muzerelle et al., 2016; Ren et al., 2018; Vertes & Linley, 2008). Previous work from our laboratory demonstrated that subsets of raphe VGLUT3 neurons project onto distinct hippocampal subregions (dHP vs vHP) (Fortin-Houde et al., 2023). Although that study raised the possibility of collateralization, it did not directly compare LS- versus vHP-projecting populations. Here, we show that while the two populations are largely segregated, a substantial proportion of neurons co-project to both targets (∼18% within VGLUT3^LS^, ∼25% within VGLUT3^vHP^). This suggests that raphe VGLUT3 projections may consist of two organizational aspects: a primary, target-specific system, and a smaller subsystem coordinating LS-vHP interactions. The identification of this architecture provides new insight into how raphe output could simultaneously influence septal and hippocampal processing in affective behaviors.

The topographical organization of VGLUT3^+^ inputs identified here may contribute to the functional specialization of LS subdivisions. Distinct VGLUT3^+^ pathways arising from raphe/IP, BNST, and NI/PCG preferentially innervate distinct LS territories, suggesting that anatomically segregated glutamatergic inputs converge onto distinct septal microcircuits. This organization provides a potential anatomical basis for the diverse functions attributed to dorsal, intermediate and ventral LS regions.

## Acknowledgments

We thank Daniel Blanchette, Véronique Pelletier and the animal facility team from the Centre Hospitalier Universitaire Sainte-Justine for assistance throughout the project. We thank Dr. Elke Küster-Schöck and Dr. Vanesa Jimenez Amilburu of the Imaging Center for their guidance and technical support in microscopy. We thank Camille Pons and Charlène Manti for their technical support for this study. Finally, we thank all members of the Amilhon laboratory for their valuable advice and constructive feedback throughout the project.

## Contributions

L.I.E. performed all experiments, analyzed data and performed statistical analysis. S.v.d.V. contributed to VGLUT3/5-HT immunohistochemistry and hippocampus/LS retrograde tracing experiments. J.F.H. contributed to the development and optimization of AAV tracing and anatomical analysis approaches. G.D. contributed to data analysis pipelines and provided programming support. L.I.E. and B.A. prepared figures and wrote the manuscript with contribution and input from the other authors. B.A. conceived and supervised the project and secured funding.

## Abbreviations

5-HT: Serotonin
B9: Supralemniscal cell group
BNST: Bed nucleus of the stria terminalis
CA1 / CA3: Cornu Ammonis hippocampal fields 1 and 3
ChAT: Choline acetyltransferase
dHP: Dorsal hippocampus
dLS: Dorsal lateral septum
DR: Dorsal raphe nucleus
DRd / DRi / DRc: Dorsal, interfascicular, and caudal subdivisions of the dorsal raphe
eYFP: Enhanced yellow fluorescent protein
GABA: Gamma-aminobutyric acid
GFP: Green fluorescent protein
HP: Hippocampus
IP: Interpeduncular nucleus
iLS: Intermediate lateral septum
LS: Lateral septum
mCherry: Monomeric red fluorescent protein
MnR: Median raphe nucleus
MRR: Median raphe region
MS: Medial septum
NAc: Nucleus accumbens
NI: Nucleus incertus
PCBs: Pericellular baskets
PCG: Pontine central gray
PMnR: Paramedian raphe nucleus
PnO: Pontine reticular nucleus
RFP: Red fluorescent protein
SFi: Septofimbrial nucleus
tdTomato: Tandem Dimer Tomato fluorescent protein
Tph2: Tryptophan hydroxylase 2
VDB: Diagonal Band of Broca, vertical limb
VGLUT1/2/3: Vesicular glutamate transporter type 1, 2, or 3
VGLUT3^LS^: LS-projecting VGLUT3+ neurons
VGLUT3^vHP^: vHP-projecting VGLUT3+ neurons
vHP: Ventral hippocampus
vLS: Ventral subdivision of the lateral septum

## References

Albert, D., & Wong, R. (1978). Hyperreactivity, muricide, and intraspecific aggression in the rat produced by infusion of local anesthetic into the lateral septum or surroundng areas. Journal of Comparative and Physiological Psychology, 92(6), 1062.

Alonso, A., Merchan, P., Sandoval, J. E., Sanchez-Arrones, L., Garcia-Cazorla, A., Artuch, R., Ferran, J. L., Martinez-de-la-Torre, M., & Puelles, L. (2013). Development of the serotonergic cells in murine raphe nuclei and their relations with rhombomeric domains. In Brain Structure & Function (Vol. 218, pp. 1229–1277). 10.1007/s00429-012-0456-8

Amilhon, B., Lepicard, E., Renoir, T., Mongeau, R., Popa, D., Poirel, O., Miot, S., Gras, C., Gardier, A. M., Gallego, J., Hamon, M., Lanfumey, L., Gasnier, B., Giros, B., & El Mestikawy, S. (2010). VGLUT3 (vesicular glutamate transporter type 3) contribution to the regulation of serotonergic transmission and anxiety. In Journal of Neuroscience (Vol. 30, pp. 2198–2210). 10.1523/JNEUROSCI.5196-09.2010

Besnard, A., Gao, Y., TaeWoo Kim, M., Twarkowski, H., Reed, A. K., Langberg, T., Feng, W., Xu, X., Saur, D., & Zweifel, L. S. (2019). Dorsolateral septum somatostatin interneurons gate mobility to calibrate context-specific behavioral fear responses. Nature Neuroscience, 22(3), 436–446.

Besnard, A., & Leroy, F. (2022). Top-down regulation of motivated behaviors via lateral septum sub-circuits. In Molecular Psychiatry (Vol. 27, pp. 3119–3128). 10.1038/s41380-022-01599-3

Blanchard, D. C., Blanchard, R. J., Lee, E. M., & Nakamura, S. (1979). Defensive behaviors in rats following septal and septal-amygdala lesions. Journal of Comparative and Physiological Psychology, 93(2), 378.

Carus-Cadavieco, M., Gorbati, M., Ye, L., Bender, F., van der Veldt, S., Kosse, C., Börgers, C., Lee, S. Y., Ramakrishnan, C., & Hu, Y. (2017). Gamma oscillations organize top-down signalling to hypothalamus and enable food seeking. Nature, 542(7640), 232–236.

Commons, K. G. (2016). Ascending serotonin neuron diversity under two umbrellas. In Brain Structure & Function (Vol. 221, pp. 3347–3360). 10.1007/s00429-015-1176-7

Cristofari, P., Desplanque, M., Poirel, O., Hebert, A., Dumas, S., Herzog, E., Danglot, L., Geny, D., Gilles, J. F., Geeverding, A., Bolte, S., Canette, A., Trichet, M., Fabre, V., Daumas, S., Pietrancosta, N., El Mestikawy, S., & Bernard, V. (2022). Nanoscopic distribution of VAChT and VGLUT3 in striatal cholinergic varicosities suggests colocalization and segregation of the two transporters in synaptic vesicles. Frontiers in Molecular Neuroscience, 15, 991732. 10.3389/fnmol.2022.991732

de Almeida, C., Chabbah, N., Eyraud, C., Fasano, C., Bernard, V., Pietrancosta, N., Fabre, V., El Mestikawy, S., & Daumas, S. (2023). Absence of VGLUT3 Expression Leads to Impaired Fear Memory in Mice. eNeuro, 10(2). 10.1523/ENEURO.0304-22.2023

Dong, H. W., & Swanson, L. W. (2004). Projections from bed nuclei of the stria terminalis, posterior division: implications for cerebral hemisphere regulation of defensive and reproductive behaviors. Journal of Comparative Neurology, 471(4), 396–433. 10.1002/cne.20002

Favier, M., Pietrancosta, N., El Mestikawy, S., & Gangarossa, G. (2021). Leveraging VGLUT3 Functions to Untangle Brain Dysfunctions. Trends in Pharmacological Sciences, 42(6), 475–490. 10.1016/j.tips.2021.03.003

Fortin-Houde, J., Henderson, F., Dumas, S., Ducharme, G., & Amilhon, B. (2023). Parallel streams of raphe VGLUT3-positive inputs target the dorsal and ventral hippocampus in each hemisphere. Journal of Comparative Neurology, 531(7), 702–719. 10.1002/cne.25452

Frahm, S., Antolin-Fontes, B., Görlich, A., Zander, J.-F., Ahnert-Hilger, G., & Ibanez-Tallon, I. (2015). An essential role of acetylcholine-glutamate synergy at habenular synapses in nicotine dependence. Elife, 4, e11396.

Goto, M., Swanson, L. W., & Canteras, N. S. (2001). Connections of the nucleus incertus. Journal of Comparative Neurology, 438(1), 86–122. 10.1002/cne.1303

Gras, C., Amilhon, B., Lepicard, E. M., Poirel, O., Vinatier, J., Herbin, M., Dumas, S., Tzavara, E. T., Wade, M. R., & Nomikos, G. G. (2008). The vesicular glutamate transporter VGLUT3 synergizes striatal acetylcholine tone. Nature Neuroscience, 11(3), 292–300.

Gras, C., Herzog, E., Bellenchi, G. C., Bernard, V., Ravassard, P., Pohl, M., Gasnier, B., Giros, B., & El Mestikawy, S. (2002). A third vesicular glutamate transporter expressed by cholinergic and serotoninergic neurons. Journal of Neuroscience, 22(13), 5442–5451. 10.1523/JNEUROSCI.22-13-05442.2002

Hale, M. W., & Lowry, C. A. (2011). Functional topography of midbrain and pontine serotonergic systems: implications for synaptic regulation of serotonergic circuits. Psychopharmacology, 213(2-3), 243–264. 10.1007/s00213-010-2089-z

Henderson, F., Dumas, S., Gangarossa, G., Bernard, V., Pujol, M., Poirel, O., Pietrancosta, N., El Mestikawy, S., Daumas, S., & Fabre, V. (2024). Regulation of stress-induced sleep perturbations by dorsal raphe VGLUT3 neurons in male mice. Cell Reports, 43(7), 114411. 10.1016/j.celrep.2024.114411

Herzog, E., Gilchrist, J., Gras, C., Muzerelle, A., Ravassard, P., Giros, B., Gaspar, P., & El Mestikawy, S. (2004). Localization of VGLUT3, the vesicular glutamate transporter type 3, in the rat brain. In Neuroscience (Vol. 123, pp. 983–1002). 10.1016/j.neuroscience.2003.10.039

Hsu, Y.-W. A., Tempest, L., Quina, L. A., Wei, A. D., Zeng, H., & Turner, E. E. (2013). Medial habenula output circuit mediated by α5 nicotinic receptor-expressing GABAergic neurons in the interpeduncular nucleus. Journal of Neuroscience, 33(46), 18022–18035.

Katrukha, E. (2020). ekatrukha/ComDet: ComDet 0.5. 3. *Zenodo*.

Kawai, H., Bouchekioua, Y., Nishitani, N., Niitani, K., Izumi, S., Morishita, H., Andoh, C., Nagai, Y., Koda, M., Hagiwara, M., Toda, K., Shirakawa, H., Nagayasu, K., Ohmura, Y., Kondo, M., Kaneda, K., Yoshioka, M., & Kaneko, S. (2022). Median raphe serotonergic neurons projecting to the interpeduncular nucleus control preference and aversion. In Nature Communications (Vol. 13, pp. 7708). 10.1038/s41467-022-35346-7

Kjelstrup, K. G., Tuvnes, F. A., Steffenach, H.-A., Murison, R., Moser, E. I., & Moser, M.-B. (2002). Reduced fear expression after lesions of the ventral hippocampus. Proceedings of the National Academy of Sciences, 99(16), 10825–10830.

Kosugi, K., Yoshida, K., Suzuki, T., Kobayashi, K., Yoshida, K., Mimura, M., & Tanaka, K. F. (2021). Activation of ventral CA1 hippocampal neurons projecting to the lateral septum during feeding. Hippocampus, 31(3), 294–304.

Leutgeb, S., & Mizumori, S. (2002). Context-specific spatial representations by lateral septal cells. Neuroscience, 112(3), 655–663.

Lima, L. B., Bueno, D., Leite, F., Souza, S., Gonçalves, L., Furigo, I. C., Donato Jr, J., & Metzger, M. (2017). Afferent and efferent connections of the interpeduncular nucleus with special reference to circuits involving the habenula and raphe nuclei. Journal of Comparative Neurology, 525(10), 2411–2442.

Liu, Y.-J., Wang, Y., Wu, J.-W., Zhou, J., Song, B.-L., Jiang, Y., & Li, L.-F. (2024). GABAergic synapses from the ventral lateral septum to the paraventricular nucleus of hypothalamus modulate anxiety. Frontiers in Neuroscience, 18, 1337207.

Lowry, C. A., Johnson, P. L., Hay-Schmidt, A., Mikkelsen, J., & Shekhar, A. (2005). Modulation of anxiety circuits by serotonergic systems. In Stress (Vol. 8, pp. 233–246). 10.1080/10253890500492787

Maita, I., Bazer, A., Blackford, J. U., & Samuels, B. A. (2021). Functional anatomy of the bed nucleus of the stria terminalis-hypothalamus neural circuitry: Implications for valence surveillance, addiction, feeding, and social behaviors. Handbook of Clinical Neurology, 179, 403–418. 10.1016/B978-0-12-819975-6.00026-1

Menon, R., Suss, T., Oliveira, V. E. M., Neumann, I. D., & Bludau, A. (2022). Neurobiology of the lateral septum: regulation of social behavior. Trends in Neurosciences, 45(1), 27– 40. 10.1016/j.tins.2021.10.010

Muzerelle, A., Scotto-Lomassese, S., Bernard, J. F., Soiza-Reilly, M., & Gaspar, P. (2016). Conditional anterograde tracing reveals distinct targeting of individual serotonin cell groups (B5–B9) to the forebrain and brainstem. Brain Structure and Function, 221(1), 535–561.

Okaty, B. W., Freret, M. E., Rood, B. D., Brust, R. D., Hennessy, M. L., deBairos, D., Kim, J. C., Cook, M. N., & Dymecki, S. M. (2015). Multi-Scale Molecular Deconstruction of the Serotonin Neuron System. Neuron, 88(4), 774–791. 10.1016/j.neuron.2015.10.007

Patel, H. (2022). The role of the lateral septum in neuropsychiatric disease. Journal of Neuroscience Research, 100(7), 1422–1437. 10.1002/jnr.25052

Paxinos, G., & Franklin, K. B. (2019). Paxinos and Franklin’s the mouse brain in stereotaxic coordinates. Academic press.

Pelosi, B., Migliarini, S., Pacini, G., Pratelli, M., & Pasqualetti, M. (2014). Generation of Pet1210-Cre transgenic mouse line reveals non-serotonergic expression domains of Pet1 both in CNS and periphery. PLoS One, 9(8), e104318. 10.1371/journal.pone.0104318

Peng, S., Yang, X., Meng, S., Liu, F., Lv, Y., Yang, H., Kong, Y., Xie, W., & Li, M. (2024). Dual circuits originating from the ventral hippocampus independently facilitate affective empathy. Cell Reports, 43(6).

Pujol, M., Fabre, V., & Daumas, S. (2023). VGLUT3 comme marqueur de vulnérabilité au stress et maladies psychiatriques associées. Médecine du Sommeil, 20(1), 20–21. 10.1016/j.msom.2023.01.003

Rashid, M., Thomas, S., Isaac, J., Karkare, S. C., Klein, H., & Murugan, M. (2025). A ventral hippocampal-lateral septum pathway regulates social novelty preference. Elife, 13, RP97259.

Ren, J., Friedmann, D., Xiong, J., Liu, C. D., Ferguson, B. R., Weerakkody, T., DeLoach, K. E., Ran, C., Pun, A., & Sun, Y. (2018). Anatomically defined and functionally distinct dorsal raphe serotonin sub-systems. Cell, 175(2), 472–487. e420.

Ren, J., Isakova, A., Friedmann, D., Zeng, J., Grutzner, S. M., Pun, A., Zhao, G. Q., Kolluru, S. S., Wang, R., Lin, R., Li, P., Li, A., Raymond, J. L., Luo, Q., Luo, M., Quake, S. R., & Luo, L. (2019). Single-cell transcriptomes and whole-brain projections of serotonin neurons in the mouse dorsal and median raphe nuclei. In Elife (Vol. 8). 10.7554/eLife.49424

Riedel, A., Stober, F., Richter, K., Fischer, K. D., Miettinen, R., & Budinger, E. (2013). VGLUT3-immunoreactive afferents of the lateral septum: ultrastructural evidence for a modulatory role of glutamate. Brain Structure & Function, 218(1), 295–301. 10.1007/s00429-012-0395-4

Riedel, A., Westerholz, S., Braun, K., Edwards, R. H., Arendt, T., & Hartig, W. (2008). Vesicular glutamate transporter 3-immunoreactive pericellular baskets ensheath a distinct population of neurons in the lateral septum. In Journal of Chemical Neuroanatomy (Vol. 36, pp. 177–190). 10.1016/j.jchemneu.2008.06.003

Risold, P., & Swanson, L. (1997). Connections of the rat lateral septal complex. Brain Research Reviews, 24(2-3), 115–195.

Risold, P. Y., & Swanson, L. W. (1997). Chemoarchitecture of the rat lateral septal nucleus. In Brain Research: Brain Research Reviews (Vol. 24, pp. 91–113). 10.1016/s0165-0173(97)00008-8

Rizzi-Wise, C. A., & Wang, D. V. (2021). Putting Together Pieces of the Lateral Septum: Multifaceted Functions and Its Neural Pathways. eNeuro, 8(6). 10.1523/ENEURO.0315-21.2021

Ryan, P. J., Ma, S., Olucha-Bordonau, F. E., & Gundlach, A. L. (2011). Nucleus incertus--an emerging modulatory role in arousal, stress and memory. In Neurosci Biobehav Rev (Vol. 35, pp. 1326–1341). 10.1016/j.neubiorev.2011.02.004

Schafer, M. K., Varoqui, H., Defamie, N., Weihe, E., & Erickson, J. D. (2002). Molecular cloning and functional identification of mouse vesicular glutamate transporter 3 and its expression in subsets of novel excitatory neurons. J Biol Chem, 277(52), 50734–50748. 10.1074/jbc.M206738200

Schindelin, J., Arganda-Carreras, I., Frise, E., Kaynig, V., Longair, M., Pietzsch, T., Preibisch, S., Rueden, C., Saalfeld, S., & Schmid, B. (2012). Fiji: an open-source platform for biological-image analysis. Nature Methods, 9(7), 676–682.

Senft, R. A., & Dymecki, S. M. (2021). Neuronal pericellular baskets: neurotransmitter convergence and regulation of network excitability. Trends in Neurosciences, 44(11), 915–924. 10.1016/j.tins.2021.08.006

Senft, R. A., Freret, M. E., Sturrock, N., & Dymecki, S. M. (2021). Neurochemically and Hodologically Distinct Ascending VGLUT3 versus Serotonin Subsystems Comprise the r2-Pet1 Median Raphe. In Journal of Neuroscience (Vol. 41, pp. 2581–2600). 10.1523/JNEUROSCI.1667-20.2021

Sheehan, T. P., Chambers, R. A., & Russell, D. S. (2004). Regulation of affect by the lateral septum: implications for neuropsychiatry. In Brain Research: Brain Research Reviews (Vol. 46, pp. 71–117). 10.1016/j.brainresrev.2004.04.009

Shutoh, F., Ina, A., Yoshida, S., Konno, J., & Hisano, S. (2008). Two distinct subtypes of serotonergic fibers classified by co-expression with vesicular glutamate transporter 3 in rat forebrain. Neuroscience Letters, 432(2), 132–136.

Sos, K. E., Mayer, M. I., Cserep, C., Takacs, F. S., Szonyi, A., Freund, T. F., & Nyiri, G. (2017). Cellular architecture and transmitter phenotypes of neurons of the mouse median raphe region. In Brain Structure & Function (Vol. 222, pp. 287–299). 10.1007/s00429-016-1217-x

Swanson, L. W., & Cowan, W. M. (1979). The connections of the septal region in the rat. Journal of Comparative Neurology, 186(4), 621–655. 10.1002/cne.901860408

Sweeney, P., & Yang, Y. (2015). An excitatory ventral hippocampus to lateral septum circuit that suppresses feeding. In Nature Communications (Vol. 6, pp. 10188). 10.1038/ncomms10188

Tingley, D., & Buzsáki, G. (2018). Transformation of a spatial map across the hippocampal-lateral septal circuit. Neuron, 98(6), 1229–1242. e1225.

van der Veldt, S., Etter, G., Mosser, C.-A., Manseau, F., & Williams, S. (2021). Conjunctive spatial and self-motion codes are topographically organized in the GABAergic cells of the lateral septum. PLoS Biology, 19(8), e3001383.

Vertes, R. P., & Linley, S. B. (2008). Efferent and afferent connections of the dorsal and median raphe nuclei in the rat. In Serotonin and sleep: molecular, functional and clinical aspects (pp. 69–102). Springer.

Wang, H. L., & Morales, M. (2009). Pedunculopontine and laterodorsal tegmental nuclei contain distinct populations of cholinergic, glutamatergic and GABAergic neurons in the rat. In European Journal of Neuroscience (Vol. 29, pp. 340–358). 10.1111/j.1460-9568.2008.06576.x

Wirtshafter, H. S., & Wilson, M. A. (2019). Locomotor and Hippocampal Processing Converge in the Lateral Septum. Current Biology, 29(19), 3177–3192 e3173. 10.1016/j.cub.2019.07.089

Wirtshafter, H. S., & Wilson, M. A. (2020). Differences in reward biased spatial representations in the lateral septum and hippocampus. Elife, 9, e55252.

Wirtshafter, H. S., & Wilson, M. A. (2021). Lateral septum as a nexus for mood, motivation, and movement. Neurosci Biobehav Rev, 126, 544–559. 10.1016/j.neubiorev.2021.03.029

Xiao, C., Wei, J., Zhang, G.-w., Tao, C., Huang, J. J., Shen, L., Wickersham, I. R., Tao, H. W., & Zhang, L. I. (2023). Glutamatergic and GABAergic neurons in pontine central gray mediate opposing valence-specific behaviors through a global network. Neuron, 111(9), 1486–1503. e1487.

Yeates, D. C. M., Leavitt, D., Sujanthan, S., Khan, N., Alushaj, D., Lee, A. C. H., & Ito, R. (2022). Parallel ventral hippocampus-lateral septum pathways differentially regulate approach-avoidance conflict. In Nature Communications (Vol. 13, pp. 3349). 10.1038/s41467-022-31082-0

